# TLR7 activation at epithelial barriers promotes emergency myelopoiesis and lung anti-viral immunity

**DOI:** 10.1101/2023.02.06.527328

**Authors:** William D Jackson, Chiara Giacomassi, Sophie Ward, Amber Owen, Tiago C. Luis, Sarah Spear, Kevin J Woollard, Cecilia Johansson, Jessica Strid, Marina Botto

## Abstract

Monocytes are heterogeneous innate effector leukocytes generated in the bone marrow and released into circulation in a CCR2-dependent manner. During infection or inflammation myelopoiesis is modulated to rapidly meet demand for more effector cells. Danger signals from peripheral tissues can influence this process. Herein we demonstrate that repetitive TLR7 stimulation via the epithelial barriers drove a potent emergency bone marrow monocyte response. This process was unique to TLR7 activation and occurred independently of the canonical CCR2 and CX3CR1 axes or prototypical cytokines. The monocytes egressing the bone marrow had an immature Ly6C-high profile and differentiated into vascular Ly6C-low monocytes and tissue macrophages in multiple organs. They displayed a blunted cytokine response to further TLR7 stimulation and reduced lung viral load after RSV and influenza virus infection. These data provide insights into the emergency myelopoiesis likely to occur in response to the encounter of single-stranded RNA viruses at barrier sites.

## Introduction

Monocytes are circulating, short-lived mononuclear phagocytes critical for host defence against infection. In mice, there are at least two subpopulations of blood monocytes defined by their expression of lymphocyte antigen 6C (Ly6C) (Geissmann et al., 2003). Ly6C-high monocytes are ‘classical’ inflammatory monocytes, which express high levels of CCR2, low levels of CX3CR1 and respond to canonical bacterial cues such as lipopolysaccharide (LPS). Ly6C-low monocytes are defined as ‘non-classical’; expressing low levels of CCR2 and high levels of CX3CR1 (Geissmann et al., 2003). These non-classical monocytes patrol the vascular lumen during times of homeostasis, surveying its integrity, orchestrating the disposal of damaged endothelial cells and subsequent inflammatory response (Auffray et al., 2007; Carlin et al., 2013; Turner-Stokes et al., 2020). This patrolling behaviour is independent of the normal leukocyte adhesion cascade and requires firm adhesion via the β2-integrin LFA1 (Carlin et al., 2013). Non-classical monocytes can respond directly to viral cues via toll-like receptor7 (TLR7), yet respond poorly to LPS (Cros et al., 2010). The existence of a heterogeneous, MHC-II-high intermediate population has also been suggested (Menezes et al., 2016; Mildner et al., 2017), but in mice the functional distinction of intermediate monocytes remains unclear. Despite widespread usage, this classification system may over-simplify monocyte heterogeneity, as mass cytometry has identified up to 8 subpopulations of monocytes in healthy human blood (Hamers et al., 2019) and novel subpopulations can emerge during inflammation and fibrosis (Satoh et al., 2017), such as Sca-1 positive ‘emergency monocytes’ during parasite infection (Abidin et al., 2017; Askenase et al., 2015).

In steady-state conditions, monocytes are produced in the bone marrow (BM) where they are derived from commitment of haematopoietic stem and progenitor cells (HSPCs) along a defined pathway that culminates in terminally differentiated mature monocytes. Although still debated, the most widely accepted of these pathways is the sequential commitment of HSPCs to the common myeloid progenitor (CMP), the monocyte-dendritic cell progenitor (MDP) and the common monocyte progenitor (cMoP), before finally generating bone marrow monocytes (Wolf et al., 2019) in a process that is dependent on the colony stimulating factor 1 (CSF1) and the transcription factor PU.1 (DeKoter et al., 1998). In addition, the work of Goodridge and colleagues suggests a functionally distinct monocyte population that can be derived from the granulocyte-monocyte progenitor (GMP), bypassing the MDP and the cMoP (Yáñez et al., 2017). The first monocyte population to be produced in the BM are the Ly6C-high monocytes, which during homeostasis are obligate precursors for Ly6C-low monocytes in a CEBP-β dependent process (Mildner et al., 2017). This is consistent with previous findings using transgenic fate mapping mice (Yona et al., 2013), direct adoptive transfer of Ly6C+ monocytes (Varol et al., 2007; Yona et al., 2013) and re-population kinetic studies following depletion regimes (Sunderkötter et al., 2004). The same process has also been suggested to occur in humans (Patel et al., 2017). Ly6C-high monocytes have a circulating half-life of <1 day in both mice and humans before converting through an intermediate stage into Ly6C-low monocytes, which can remain in the blood for between ∼2 and 7 days (Patel et al., 2017; Yona et al., 2013). The egression of Ly6C-high monocytes from the BM is largely dependent on the C-C chemokine receptor type 2 (CCR2) (Tsou et al., 2007). Similarly, Ly6C-high monocyte extravasation into tissues is mediated by CCR2 in response to local production of the C–C motif chemokine ligand 2 (CCL2) or 7 (CCL7) (Shi and Pamer, 2011). However, the dynamics of monocyte subpopulation production and BM egression during inflammation and/or infection remain poorly understood.

During infection or tissue injury monocytes are crucial for controlling the invading pathogen (Haist et al., 2017; Shi and Pamer, 2011) or for regulating the tissue repair process (Wynn and Vannella, 2016). For example, the recruitment of Ly6C-high inflammatory monocytes in the lungs in response to respiratory pathogens like respiratory syncytial virus (RSV) is essential to control viral load and lessen disease severity (Goritzka et al., 2015). As such, the process of myelopoiesis is differentially regulated during infection or inflammation to rapidly meet demand for ‘emergency’ effector monocytes or neutrophils. An acute requirement for additional monocytes can either be met by the spleen via a reservoir of mature splenic monocytes and accompanying extramedullary haematopoiesis (Swirski et al., 2009), or by conventional BM haematopoiesis (Wolf et al., 2019). A key component of the ‘emergency’ process is sensing and communicating the danger signals to the haemopoietic progenitor pool in either the BM or the spleen. This occurs primarily through activation of pattern recognition receptors (PRRs), most notably the TLR family, whose expression has been confirmed on HSPCs in both human and mouse (Nagai et al., 2006; Sioud et al., 2006). The sensing of pathogens by the haematopoietic system can either occur directly or as a secondary effect of inflammatory mediators produced at the barrier sites such as the skin or the gut (Askenase et al., 2015; Baldridge et al., 2010; Pietras et al., 2016) or by BM stromal cells (Shi and Pamer, 2011). The nature of the distal signals from barrier sites to BM and the features of the ‘emergency’ processes triggered under different settings remain largely unknown.

The skin is the largest barrier site in the body and is targeted as an entry point by a variety of pathogens, perhaps most notably the diverse group of arboviruses spread by mosquitoes. These infect hundreds of millions of people annually and include serious threats to human health such as dengue virus and Zika virus, both of which are single-stranded RNA viruses sensed by TLR7 (Paixão et al., 2018). Recently, it has been shown that the skin immune response via TLR7 expressed in dermal dendritic cells can locally protect against a second viral infection at the inoculation site (Bryden et al., 2020). However, it is unclear what effect TLR7 activation at the epithelial barrier has on the haematopoietic system and the subsequent innate immune response in other organs. Here we used R848, a TLR7/8 agonist, and demonstrate that only persistent TLR7 stimulation at an epithelial barrier such as the skin or gut was able to drive a unique CCR2-independent BM emergency monocyte response. This process was characterised by the release of immature Ly6C-high pre-monocytes into the periphery and their differentiation to both Ly6C-low monocytes in the blood and to tissue macrophages in multiple organs. The emergency monocytes released by the BM under these conditions displayed an impaired response to TLR7 restimulation and promoted lung viral control, which dampened disease severity after RSV and influenza virus infection.

## Results

### Persistent TLR7 stimulation at epithelial barrier sites drives systemic monocytosis

To investigate how the hematopoietic response to a single-stranded RNA virus could be influenced by the point of entry, we administered a TLR7 agonist, R848, to BALB/c mice topically or intraperitoneally (I.P) (100µg, 3 times a week for 4 weeks, equivalent to 12 treatments) (**Fig S1A**) and observed a marked monocytosis only in the topical R848 group (**Fig 1A**). While blood monocyte counts (CD11b+CD115+) in vehicle and I.P treated mice remained largely unchanged (mean of 850.8±159/ µl), mice that received topical R848 had a mean monocyte count of 25,154±2,956/ µl, an approximately 30-fold expansion (**Fig 1A**). The monocytosis was accompanied by a ratio switch in the Ly6C-high and Ly6C-low monocyte subpopulations, with the Ly6C-low population accounting for >90% of total monocytes after topical R848 (**Fig 1A**). We also found a pronounced splenomegaly in the topically treated mice which was not present in the mice receiving R848 I.P (**Fig S1B**). To investigate the kinetics of the monocytosis mice were treated with topical R848 4 times and blood samples were taken 24 hours after each treatment. After 1 treatment, we observed a marked drop in monocyte (**Fig 1B**) and lymphocyte (both CD3+ T-cell and B220+ B-cell) counts (**Fig S1C**). While the lymphocyte counts slowly returned within the normal range by 4 treatments (**Fig S1C**), both monocyte subpopulations expanded quickly after 2 treatments (**Fig 1B**). On subsequent R848 applications the Ly6C-low monocyte population continued to expand, whilst the level of Ly6C-high monocytes remained stable (**Fig 1B**), ultimately leading to a ratio switch (**Fig 1C**) as in **Fig 1A**. Throughout the time course no change was seen in blood neutrophil counts (CD11b+Ly6G+Ly6C-low), indicating that topical R848 does not trigger a pan-myeloid response (**Fig 1D**). To confirm that the monocyte response was not due to the systemic diffusion of the compound, we administered R848 intravenously (I.V) and compared it to topical application. While both routes of administration caused lymphopenia, only the topical route was able to promote monocytosis and a ratio switch towards Ly6C-low monocytes (**Fig 1E**). We then reasoned that perhaps the monocyte response triggered by TLR7 activation was specific to epithelial barrier sites. To test if this was a unique response to topical challenge, R848 was dissolved in the drinking water at a concentration that was calculated to be equivalent to the dose applied topically (Bachmanov et al., 2002). Oral R848 induced a monocytosis with a proportional switch towards Ly6C-low monocytes that was comparable to that seen after topical treatment (**Fig 1F**).

**Figure 1:**
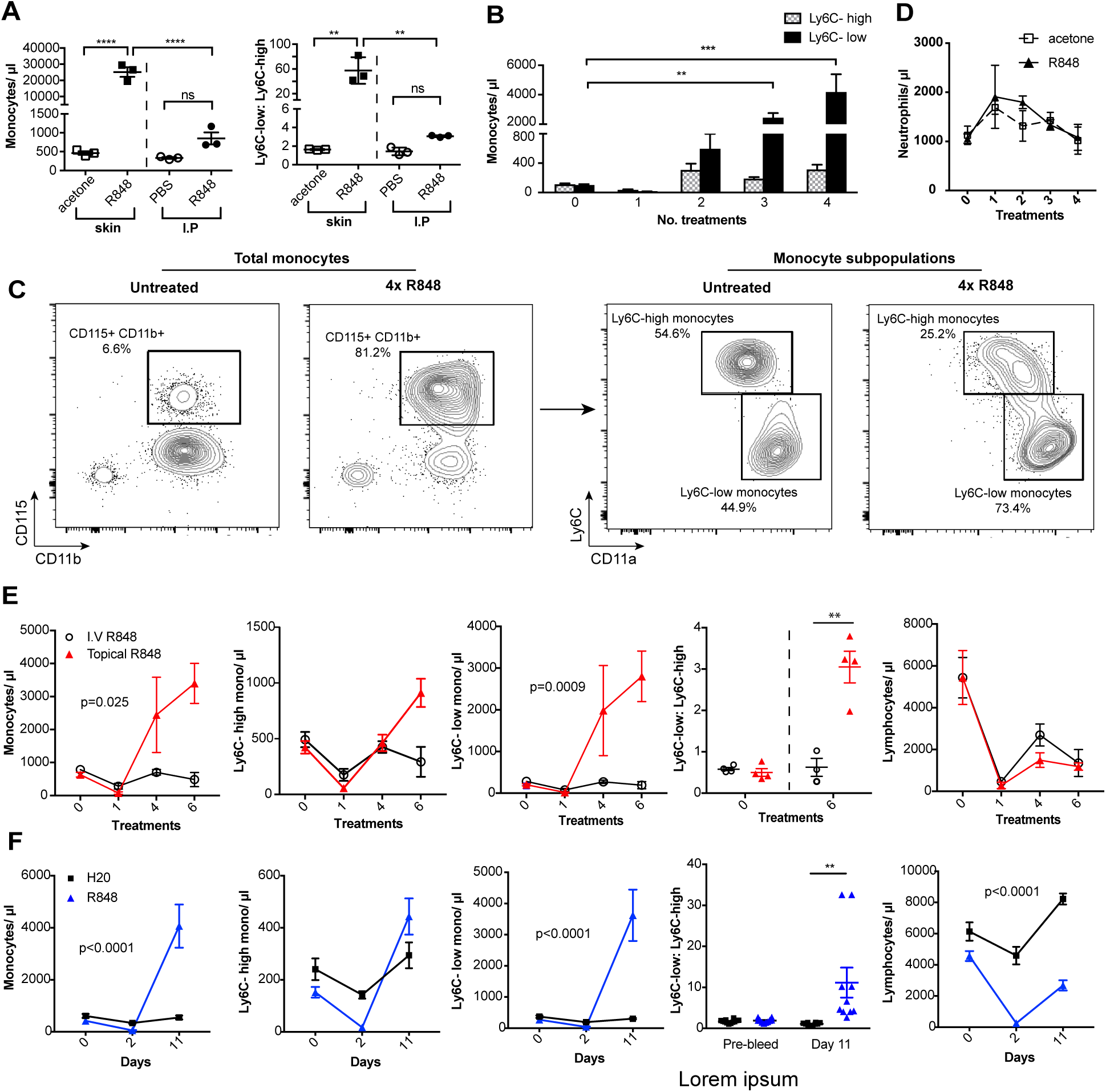
Repetitive R848 administration at barrier sites drives a profound monocytosis. **A)** BALB/c mice (n=3 per group) received 100µg of R848 topically or I.P, 3x per week for 4 weeks. Control BALB/c mice were given topical acetone or 200µl of PBS I.P. Left panel shows the number of total monocytes (CD11b+CD115+), the right panel the ratio between Ly6C-high and Ly6C-low monocytes. **B)** BALB/c mice (n=3 per group) received 4 treatments of topical R848 (100µg). Numbers of Ly6C-high (grey bars) and Ly6C-low (black bars) monocyte 24 hours after each treatment. **C)** Representative plots of total monocytes (left panel) and of subpopulations (right panel) in mice treated topically with acetone or R848 4 times. **D)** Mice treated as in B. Neutrophil counts 24 hours after each treatment. **E)** BALB/c mice (n=4 per group) received 6 treatments with 100µg topical (red line) or I.V R848 (black line). Blood counts are shown for total monocytes, Ly6C-high monocytes, Ly6C-low monocytes, monocyte subpopulation ratio and lymphocytes at 24 hours after the indicated treatment. **F)** C57BL/6 mice were given drinking water containing 8.3µg/ ml R848 (blue line, n=10) or vehicle control (black line, n=12) for 11 days. Blood counts are shown for total monocytes, Ly6C-high monocytes, Ly6C-low monocytes, monocyte subpopulation ratio and lymphocytes at the indicated time point. Data representative of at least 2 independent experiments (except **A**). One-way ANOVA with Bonferroni’s multiple comparison test (**A**); two-way ANOVA with Tukey’s multiple comparison for analysis of time-course experiments (**B, D, E, F**). Data are the mean ± SEM; only significant p values are indicated; ** p<0.01; *** p<0.001; **** p<0.0001. Please also refer to Fig S1

As the response to other TLR stimuli such as LPS is dose-dependent and repeated administrations can either result in sensitization or tolerance, which is considered a form of ‘innate immune memory’ (Biswas and Lopez-Collazo, 2009), we performed a dose-response experiment with R848 given I.P. None of the doses tested caused a significant increase in blood monocyte counts when compared to vehicle (**Fig S1D**). In addition, the monocytosis following topical R848 was clearly dose dependent, with 100µg inducing a strong response, 1µg causing no change and the 10µg group displaying an intermediate phenotype (**Fig S1E**). Consistent with the dose response data, application of imiquimod (IMQ), which is a ∼100-fold less potent agonist, did not elevate peripheral monocyte counts even after 6 daily treatments (**Fig S1F**). Together these data demonstrate that a persistent and potent TLR7 stimulation at an epithelial barrier can trigger a distinctive innate immune response characterised by a profound monocytosis.

### Skin-induced monocytosis requires TLR7 activation

We next sought to investigate whether the monocyte response via an epithelial barrier could also be promoted by other TLR stimuli. We therefore treated mice topically with LPS or poly I:C (TLR4 and TLR3/ RIG-I agonists respectively), alongside an R848 control group. For this experiment we utilized a modified water-soluble version of R848 to allow all agonists to be dissolved in the same vehicle. As previously, topical R848 triggered an immediate leukopenia followed by the characteristic monocytosis dominated by Ly6C-low monocytes, whereas LPS and poly I:C had no substantial effects on blood cell counts (**Fig 2A**). Similarly, topical application of a TLR9 agonist, CpG oligodeoxynucleotides (ODNs), neither changed monocyte counts nor caused lymphopenia (**Fig 2B**). Moreover, topical application of the potent pro-inflammatory stimulus 12-O-Tetradecanoylphorbol-13-acetate (TPA) did not affect monocyte counts after 4 treatments (**Fig 2C**). In view of the marked difference between the effects induced by R848 and the other TLR agonists, we next used TLR7-deficient (*Tlr7^-/-^*) mice to exclude activation of TLR8 or inflammasome, or any off-target effects of the R848 compound. Total monocytes and Ly6C-low monocytes were significantly elevated in the wild-type (WT) animals, while *Tlr7^-/-^* mice did not show a response to R848 confirming the specificity of the pathway involved (**Fig 2D**).

**Figure 2:**
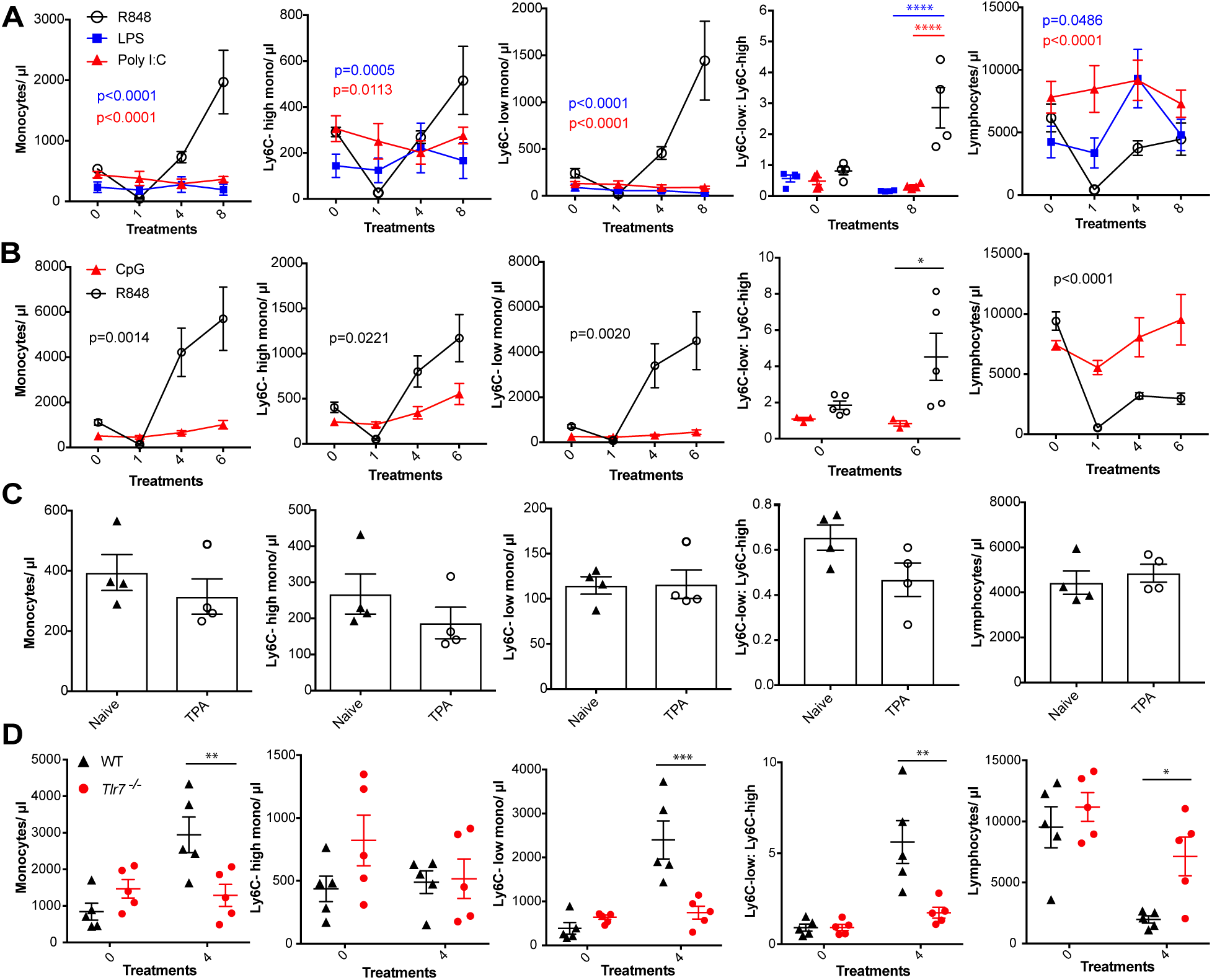
R848-induced monocytosis is specific to TLR7 activation. Blood counts are shown for total monocytes, Ly6C-high monocytes, Ly6C-low monocytes, monocyte subpopulation ratio and lymphocytes 24 hours after the indicated treatment. **A)** BALB/c mice (n=4 per group) received 8 treatments with topical R848 (100µg, black line), LPS (100µg, blue line) or Poly l:C (100µg, red line). **B)** C57BL/6 mice received 6 treatments with topical R848 (100µg, n=5, black line) or CpG (100µg, n=3, red line). ***C)*** BALB/c mice (n=4 per group) received 4 topical treatments with 2.5nmol TPA. **D)** C57BL/6 mice (n=5, black triangles) or Tlr7-/-mice (n=5, red circles) received 4 treatments with topical R848. Data representative of 2 independent experiments (except **A** and **C**). Two-way ANOVA with Tukey’s multiple comparison for time-course experiments (**A, B**); unpaired t-test (**C, D**). Data are the mean ± SEM; only significant p values are indicated; * p<0.05; ** p<0.01; *** p<0.001. Please also refer to Fig S2.

Various cytokines have previously been implicated in the regulation of myeloid cell production during inflammation, most notably the interferon (IFN) family and IL1β (Askenase et al., 2015; Buechler et al., 2013; Mitroulis et al., 2018). We found no obvious differences in monocyte counts after topical R848 treatments in either IFN-y-or IFNAR1-deficient mice compared to the WT animals ((**Fig S2A** and **Fig S2B**), discounting a role for type I and type II IFNs. We then utilized the IL-1 receptor antagonist anakinra, which has been reported to cross react with the mouse protein (Iannitti et al., 2016), and observed no differences in the induction of monocytosis or the monocyte subpopulation ratio (**Fig S2D**) making the involvement of IL-1 unlikely. In addition, blocking IL-6 and TNF-α signalling did not prevent the onset of monocytosis (**Fig S2C** and **S2E**).These data collectively suggest that only activation of the TLR7 pathway at the skin barrier can drive the changes in the myeloid compartment that occur independently from inflammasome activation or some of the well-established cytokine-mediated pathways.

### Myeloid cells orchestrate the R848-induced monocytosis

We next attempted to determine which cells initiate the monocyte response. We first generated BM chimeras using *Tlr7^-/-^* mice (Hemmi et al., 2002) to distinguish between stromal and haematopoietic cells. BM reconstituted mice were treated with topical R848 and monocyte response assessed. After 4 treatments, there was a marked increase in the proportion of total monocytes and Ly6C-low monocytes in the mice that received WT BM, but not in mice that received *Tlr7^-/-^* BM, regardless of the host genotype (**Fig S3A**), indicating that cells of haematopoietic origin and not irradiation-resistant skin-resident cells like stromal or epithelial cells were responsible for the myeloid response to R848.

To identify the BM-derived cells responding to the TLR7 activation, we first utilized *Rag2^-/-^* mice, which lack both T- and B-cells (Shinkai et al., 1992). After 4 topical treatments with R848, both WT and *Rag2^-/-^* animals developed a monocytosis dominated by Ly6C-low cells (**Fig 3A**), indicating that lymphocytes are dispensable. We then crossed *Tlr7*-floxed mice (Solmaz et al., 2019) with *Lyz2^Cre/Cre^* animals, which express the Cre recombinase in monocytes, neutrophils and some macrophage populations (Abram et al., 2014). While after R848 treatment both WT and *Lyz2^Cre/+^ x Tlr7^fl^* mice developed monocytosis, this expansion was reduced by >50% in the mice lacking TLR7 in *Lyz2*-expressing cells (**Fig 3B**), demonstrating that Lyz2-expressing cells were at least partially responsible for the skin response to TLR7 activation. As mast cells, eosinophils and basophils are not reported to express *Lyz2* (Abram et al., 2014) we investigated these cells individually. *ΔdblGATA* mice, lacking eosinophils, and *Cpa3^Cre/^*^+^ mice, lacking both mast cells and basophils, did not show any obvious defect in their response to R848 (**Fig S3B** and **S3C**). In addition, neutrophils depletion using an anti-Ly6G depletion antibody (Daley et al., 2008) failed to abolish the monocytosis. On the contrary, it increased the R848-induced monocytosis (**Fig S3D**) consistent with the observation that (Cortez-Retamozo et al., 2012; Swirski et al., 2009)neutrophil depletion can trigger a mild degree of monocytosis (Patel et al., 2019). In light of these observations, we concluded that a BM-derived myeloid population was contributing to promote the systemic response following TLR7 activation.

**Figure 3:**
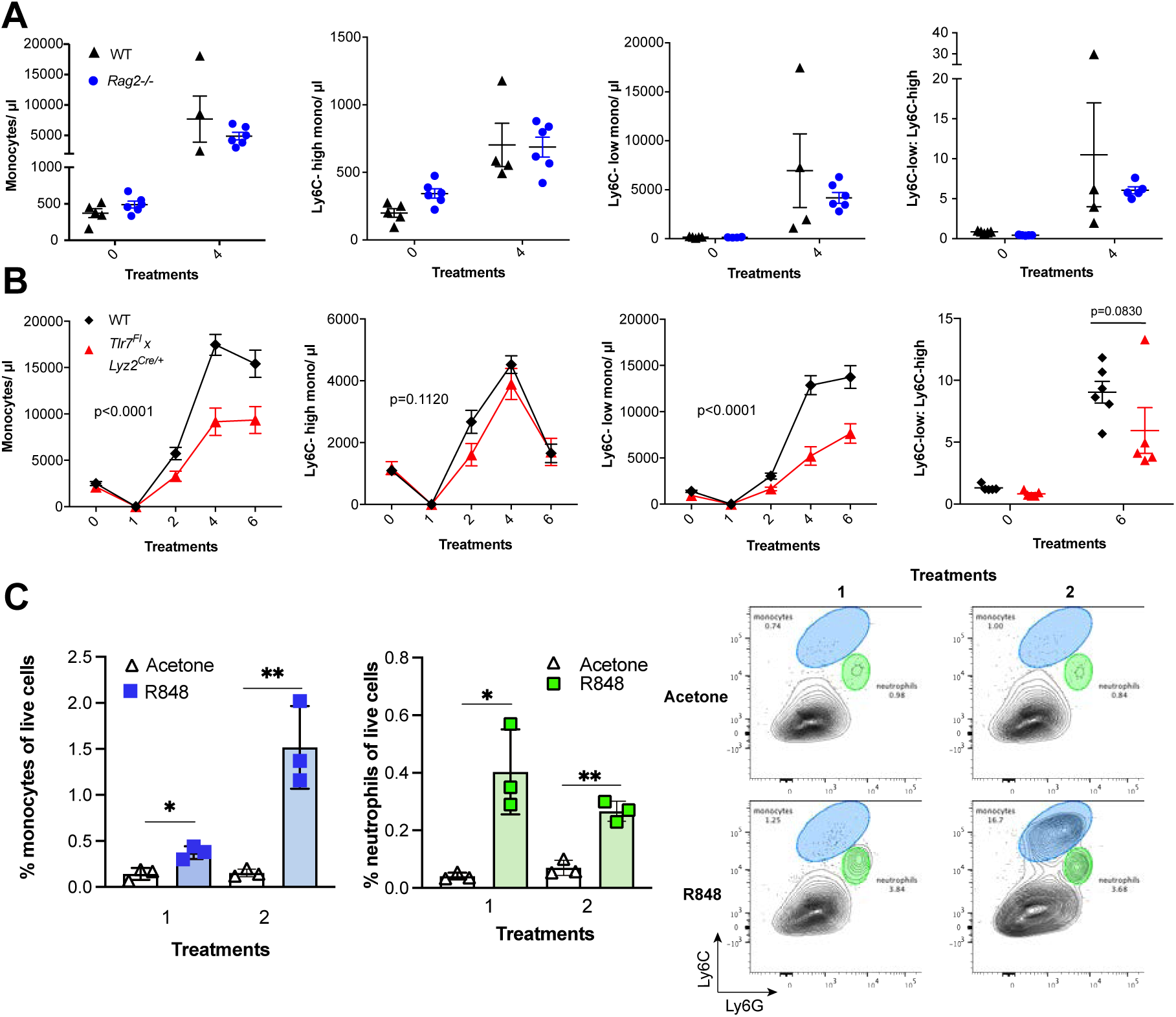
R848-induced monocytosis is driven by TLR7 activation of myeloid cells. **(A-B)** Blood counts for total monocytes, Ly6C-high monocytes, Ly6C-low monocytes, monocyte subpopulation ratio and lymphocytes at baseline and 24 hours after the indicated treatment. **A)** C57BL/6 mice (n=5, black triangles) and Rag2-/-mice (n=6, blue circles) received 4 topical treatments with R848. **B)** C57BL/6 mice (n=6, black rhombi) and Tlr7fl x Lyz2Cre/+ mice (n=5, red triangles) received 6 topical treatments with R848. **C)** C57BL/6 mice (n=3 per group) were treated once or twice with topical R848 or acetone. Treated ear skin was harvested at 24 hours post-treatment and analysed by flow cytometry. Proportion of monocytes (CD11b+Ly6C+Ly6G-low) and neutrophils (CD11b+Ly6G-high-Ly6C-low) among total live cells (left panels). Representative flow cytometry plots gated on CD11b+ cells (right panels). Data represent a single experiment (A, C) or 2 experiments (B). Two-way ANOVA, with Tukey’s multiple comparison for time-course experiments (A, B); unpaired t-test (C). Data are the mean ± SEM; only significant p values are indicated; * p<0.05; ** p<0.01. Please also refer to Fig S3.

We next explored whether local tissue infiltrating myeloid cells were involved in the systemic response. We applied R848 twice and observed infiltration of monocytes in the skin after the first treatment (**Fig 3C**) suggesting that these BM-derived myeloid cells were not only involved in triggering the initial response to TLR7 stimulation, but also acted in a positive feedback manner and played a key role in maintaining the systemic myelopoiesis.

### The emergency monocyte response triggered by cutaneous R848 originates in the BM

While haematopoiesis of myeloid cells during homeostasis occurs predominantly in the BM, under pathological conditions this can be superseded by extramedullary haematopoiesis in the spleen (Cortez-Retamozo et al., 2012; Swirski et al., 2009). To understand the source of the monocyte response triggered by cutaneous TLR7 activation we performed splenectomy or sham surgery and applied topical R848 treatment after the mice had fully recovered from the surgery. Monocytosis and a ratio switch towards Ly6C-low monocytes occurred in the R848-treated mice with or without spleen and were markedly different from the untreated splenectomised mice (**Fig 4A**) indicating that the stimulation of TLR7 at the skin barrier was driving a haematopoietic response mainly in the BM. Of note, the R848-treated splenectomised mice showed a slightly enhanced phenotype, suggesting that the spleen may retain some of the circulating monocytes. To confirm the BM origin of the R848-induced monocytosis we conducted *in vivo* fate-tracing experiments using HSC-SCL-CreER^T^;R26EYFP mice (Göthert et al., 2005). Under our experimental conditions tamoxifen administration induced recombination in ∼45% of the BM HSCs (**Fig S4A-B**) with no recombination in the peripheral blood. Consistent with previous experiments the tamoxifen-treated HSC-SCL-CreER^T^;R26EYFP mice responded to the topical R848 with a marked increase in the total monocyte count and a ratio switch towards Ly6C-low monocytes (**Fig 4B**) indicating that neither the genetic modification nor the tamoxifen had altered the BM response. Importantly, the same pattern was observed among the EYFP-positive monocytes that were predominantly Ly6c-low monocytes (**Fig 4C**) confirming that the R848-induced monocytes originates mainly from the BM.

**Figure 4:**
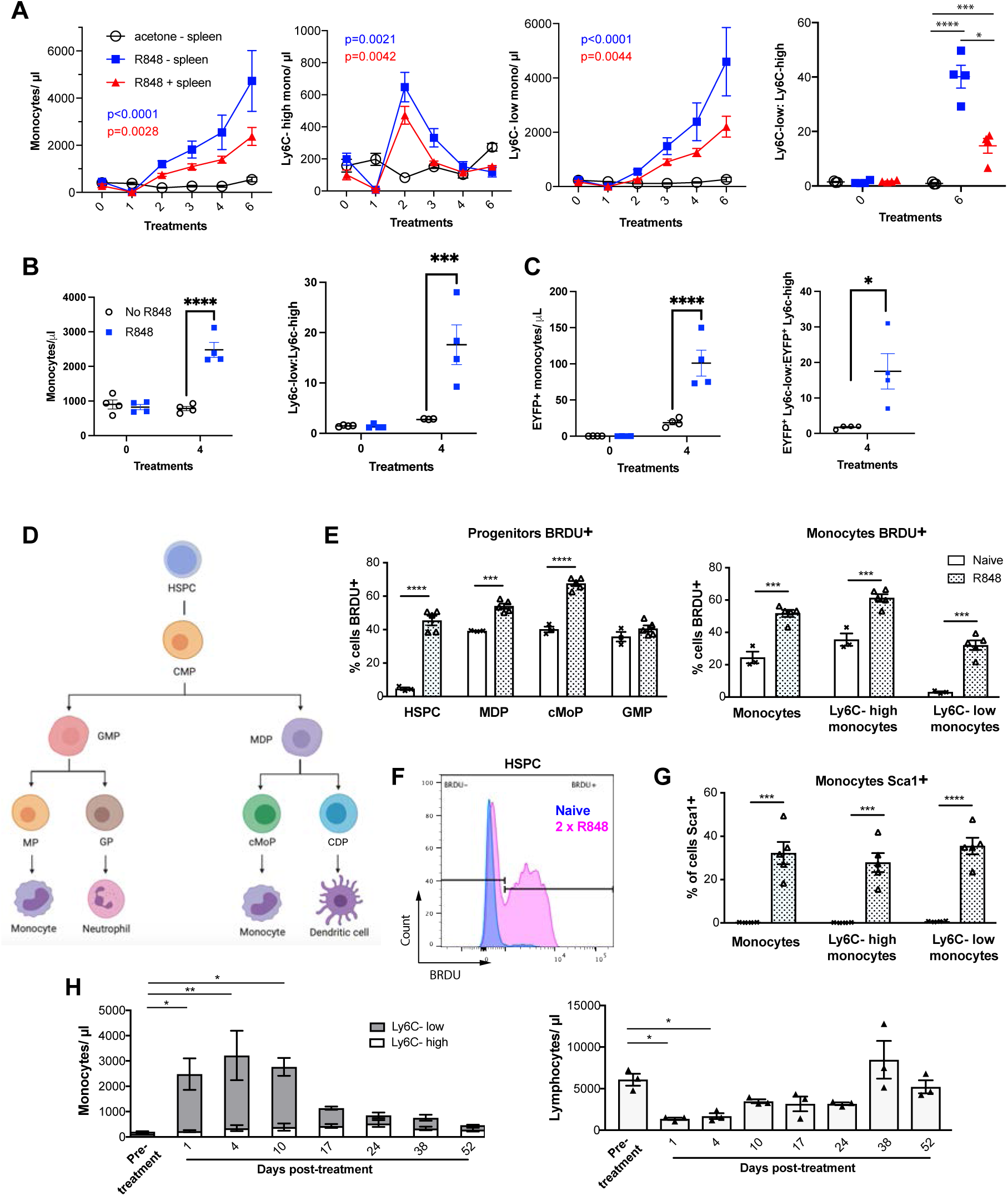
R848-induced monocytes are derived from the BM and have features of emergency myelopoiesis. **A)** BALB/c mice underwent splenectomy or sham surgery and were left to recover for 7 weeks. Among the splenectomised mice, a group was treated 6x with topical R848 (n=4, blue line) and the remaining mice with acetone (n=3, solid black line). The sham surgery group was treated with R848 (n=4, red line). Blood counts are shown for total monocytes, Ly6C-high monocytes, Ly6C-low monocytes and the monocyte subpopulation ratio at baseline and 24 hours after the last treatments. **B-C)** HSC-SCL-Cre-ERT;R26R-EYFP mice (n=4 per group) received Tamoxifen (4mg/100μL) by oral gavage for 5 consecutive days. Three days later, the left ears were treated topically 4x with R848 (blue squares) or left untreated (open circles). Shown are the total blood monocyte counts and the monocyte subpopulation ratio (**B**); the total blood EYFP+ monocyte counts and the subpopulation ratio among the EYFP-expressing monocytes at baseline and 24 hours after the last treatment (**C**). **D)** Diagram illustrating BM myeloid progenitor differentiation: haematopoetic stem and progenitor cells (HSPC); common myeloid progenitor (CMP); monocyte-dendritic precursor (MDP); granulocyte-monocyte progenitor (GMP); common monocyte progenitor (cMoP); common dendritic progenitor (CDP); monocyte-committed progenitor (MP); granulocyte-committed progenitor (GP). **(E-G)** C57BL/6 mice were injected with 2mg BRDU I.P, either naïve (n=3, white bars) or after 2 topical R848 treatments (n=5, grey bars). Mice culled at 16 hours after the BRDU injection and BM harvested. **E)** Percentage of BRDU positivity in HSPC, MDP, cMoP, GMP, total monocytes, Ly6C-high monocytes and Ly6C-low monocytes. **F)** Representative histogram of BRDU expression in HSPC at baseline (blue) or after 2x topical R848 (magenta). **G)** Percentage of Sca1 positivity in total monocytes, Ly6C-high monocytes and Ly6C-low monocytes in the BM. **H)** BALB/c mice (n=3) received 4 treatments with topical R848. Blood counts for Ly6C-high monocytes (white bars), Ly6C-low monocytes (grey bars) and lymphocytes were monitored at the indicated timepoints after the cessation of treatment. Data representative of 2 independent experiments (except **A** and **H**). Two-way ANOVA with Tukey’s multiple comparison for time-course experiments **(A-C, H)**; unpaired t-test **(E, G)**. Data are the mean ± SEM; only significant p values are indicated; * p<0.05; ** p<0.01; *** p<0.001; **** p<0.0001. Please also refer to Fig S4.

Monocytes are derived from a defined program of progenitor differentiation in the BM outlined in **Fig 4D**. To investigate this pathway, we assessed proliferation of bone marrow progenitor cells using bromodeoxyuridine (BRDU) pulse-chase experiments. When we examined BRDU incorporation in progenitor cells (gated strategy in **Fig S4C**), we found that R848 dramatically increased the percentage of HSPCs in cell cycle from ∼4.6% to ∼45.6% (**Fig 4E** and **Fig 4F**). Consistent with the peripheral blood data showing that TLR7 activation targets the monocytes and does not affect the granulocyte lineage, there was a significant increase in BRDU positivity among MDPs and cMoPs, but not in GMPs (**Fig 4E**). Unsurprisingly, the mature bone marrow monocyte population also showed increased BRDU incorporation (**Fig 4E**) and this was present in both monocyte subpopulations indicating that effects of TLR7 activation were not limited to the Ly6C-low cells (**Fig 4E**). Of note, the R848-induced bone marrow monocytes also up-regulated expression of the stem cell antigen 1 (Sca1) (**Fig 4G**), a feature consistent with an emergency myelopoiesis (Askenase et al., 2015).

Given the dramatic increase in the proportion of normally quiescent HSPCs in cell cycle, we hypothesized that R848-induced monocytosis would persist for a considerable period after cessation of the treatment. Indeed, after 4 topical R848 treatments, it took at least 17 days for total monocyte counts to return to pre-bleed levels (**Fig 4H**). The monocyte changes were accompanied by a decrease in blood lymphocyte numbers, which reached pre-treatment levels by day 10 (**Fig 4H**). We therefore concluded that cutaneous TLR7 activation can bypass the splenic reservoir and can instruct the BM HSPCs to proliferate and differentiate predominantly along the monocytic pathway triggering an emergency myelopoiesis.

### Topical R848 accelerates the differentiation of Ly6C-high monocytes to macrophages

Under homeostatic conditions, Ly6C-high monocytes have been shown to be obligate precursors for Ly6C-low monocytes in the blood (Mildner et al., 2017). As both BM monocyte subpopulations displayed increased proliferation after topical R848, we questioned whether the Ly6C-low monocytosis under the pathological setting of our experimental model was the result of increased Ly6C-high monocyte conversion or of enhanced egression of Ly6C-low monocytes emerging from the BM as a separate lineage. To answer this question, we utilized an antibody-mediated depletion strategy via injection of an Fc-chimeric mouse anti-GR1. In naïve mice this antibody fully depletes blood Ly6C-high monocytes by 24 hours post-injection, leaving Ly6C-low cells unaffected (**Fig 5A**). We then administrated this depleting antibody or an anti-FITC isotype control for 3 consecutive days to R848-treated mice, starting simultaneously with the 3^rd^ R848 treatment at day 6. As expected in untreated mice the Fc-chimeric mouse anti-GR1 antibody reduced the Ly6C-high monocyte counts to undetectable levels (**Fig 5B**). Importantly, in R848-treated mice the Ly6C-high monocyte depletion due to the Fc-chimeric mouse anti-GR1 antibody abrogated the monocytosis, whilst the isotype control had no effect (**Fig 5B**). These data suggest that the expanded Ly6C-low monocyte population is predominantly, if not entirely, derived from the Ly6C-high monocyte population and not from an expansion of a separate lineage in the BM.

**Figure 5:**
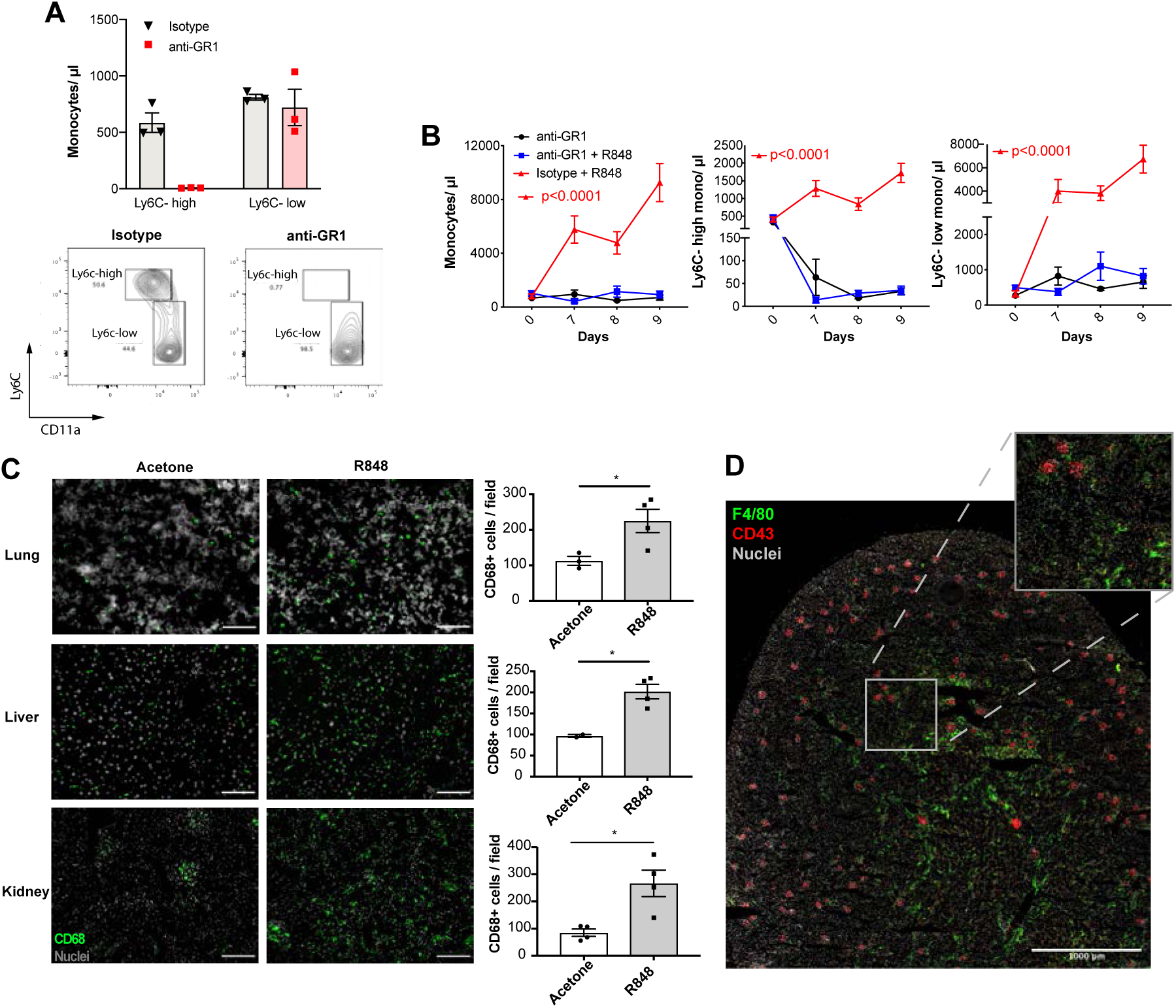
Topical R848 drives Ly6C-high monocyte differentiation to intravascular and tissue macrophages. **A)** C57BL/6 mice were injected I.P with either anti-GR1 antibody (n=3, red squares) or matched isotype control (n=3, black triangles). Blood Ly6C-high and Ly6C-low monocyte counts (upper panels) and representative flow cytometry plots (bottom panels) are shown at 24 hours post-injection. **B)** C57BL/6 mice were treated with combinations of topical R848 and I.P anti-GR1 antibody or matched isotype control, with the antibody injected daily for 3 consecutive days starting 1 hour prior to the 3rd R848 treatment (day 6). The groups were: isotype control and R848 (n=4, red triangle), anti-GR1 and R848 (n=4, blue squares), anti-GR1 alone (n=4, black circles). Blood counts for total monocytes, Ly6C-high monocytes and Ly6C-low monocytes at baseline and on the indicated days are shown. Statistics compare anti-GR1+R848 to isotype+R848. **C)** C57BL/6 mice received 4 treatments with topical R848 (n=4 per group, grey bars) or acetone (n=4 per group, white bars). Immunofluorescent staining for CD68 (green) and nuclei (grey) was performed on tissue sections from lung, liver and kidneys. Staining was quantified as mean CD68+ cells per field. **D)** C57BL/6 mice (n=4) received 6 treatments with topical R848. Kidney sections were stained for CD43 (red), F4/80 (green) and nuclei (grey). A representative tile scan is shown. Data representative of two independent experiments (except **D**). Time-course experiments analysed with two-way ANOVA with Tukey’s multiple comparison used to compare between groups at a given time point (**B**); comparison of two groups at a single timepoint calculated using unpaired t-test (**C**). Data are the mean ± SEM; only significant p values are indicated; * p<0.05.

Ly6C-low blood monocytes are viewed as intravascular ‘blood macrophages’ due to their transcriptional similarity to tissue macrophages (Mildner et al., 2017). Therefore, we reasoned that the cutaneous R848 treatment may be driving an accelerated macrophage differentiation program in Ly6C-high monocytes and thereby also promoting macrophage infiltration into tissue. To confirm this, tissue sections from lungs, liver and kidneys of R848-treated mice were stained with an anti-CD68 antibody. A significant increase of CD68+ cells per field was observed in all these organs (**Fig 5C**). In the kidney we also used a combination of CD43 and F4/80 staining to distinguish intravascular CD43-high non-classical monocytes from F4/80-high CD43-negative tissue macrophages (Kuriakose et al., 2019). Using this strategy, we found that CD43-positive cells were confined to the vasculature of the glomeruli, while F4/80-high macrophages were mainly located in the medulla (**Fig 5D**), findings consistent with the different roles of these two myeloid subpopulations.

Together these data indicate that cutaneous TLR7 activation triggers the egression from the BM of emergency Ly6C-high monocytes that undergo an accelerated macrophage differentiation, both patrolling the vascular lumen as Ly6C-low monocytes and also directly invading different organs as blood-derived tissue macrophages. As the Ly6C-low population remains in the intravascular space, this would explain the progressing switch in the Ly6C-high and Ly6C-low monocyte ratio (**Fig 1B**).

### Topical R848 triggers a CCR2-independent myeloid response

Under steady-state conditions, the monocyte release from the BM is orchestrated by CCR2 and CX3CR1 (Landsman et al., 2009; Tsou et al., 2007). We investigated whether this was also true for the R848-induced myelopoiesis using *Ccr2^RFP/RFP^* knock-in mice (Saederup et al., 2010). As previously reported (Tsou et al., 2007), at baseline CCR2-deficient mice had less blood monocytes than the WT animals due to a failure of Ly6C-high monocytes to egress from the BM (**Fig 6A**). After topical R848 treatments, monocytosis was present in both strains and the Ly6C-low monocyte fold increase from baseline was even higher in the mice lacking CCR2 (**Fig 6B**). Consistent with the notion that the R848-driven monocyte response bypassed the conventional pathways, we found a similar increase of CD68+ cells per field in the kidney and lung of the CCR2-deficient mice indicating that different signals also orchestrated the monocyte extravasation into the tissues (**Fig 6C**).

**Figure 6:**
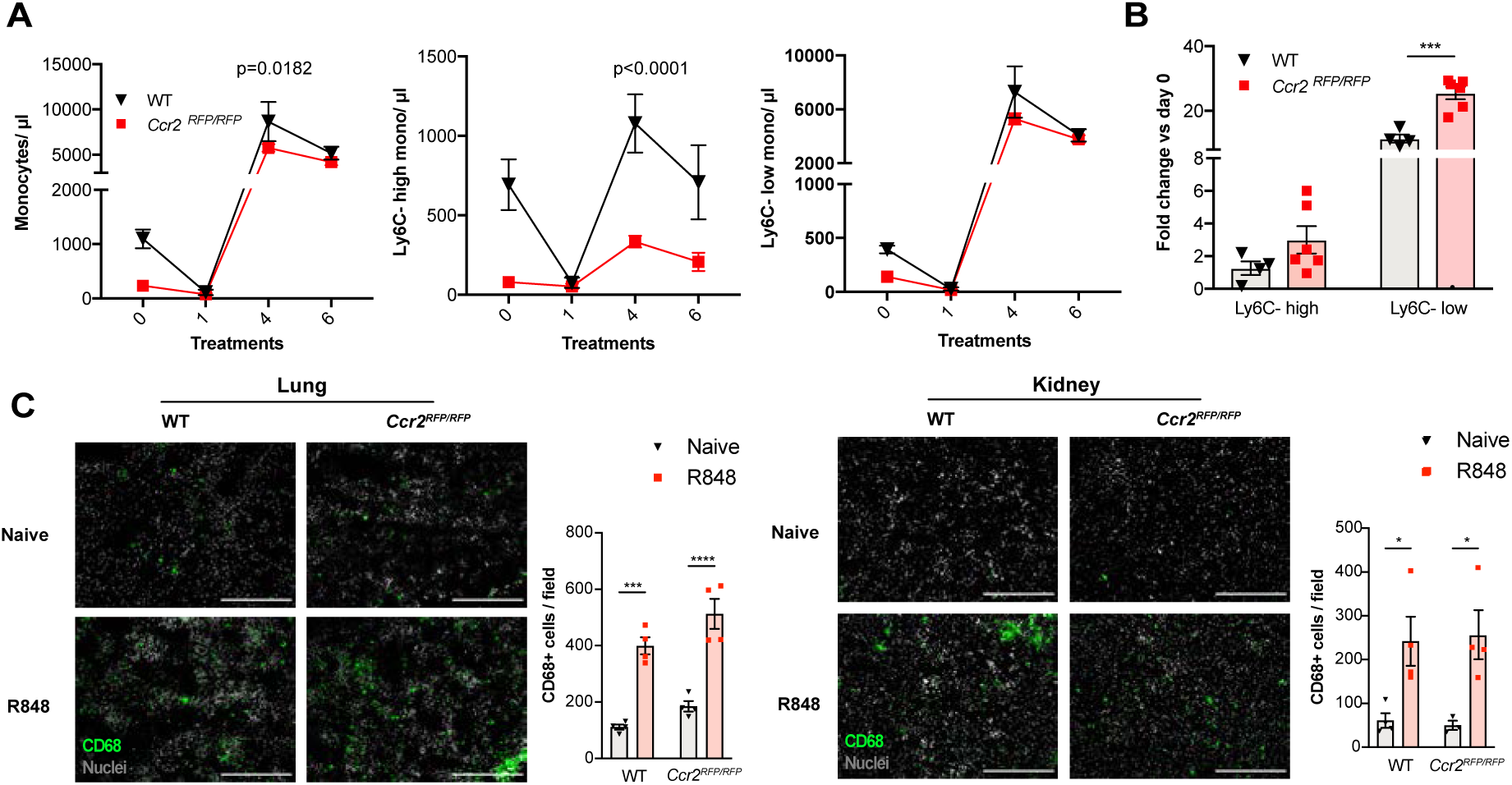
Immature monocytes egress from the BM independently of CCR2. **A-B)** C57BL/6 mice (n=4, black triangles) or Ccr2RFP/RFP mice (n=6, red squares) were treated topically 6 times with R848. **A)** Blood counts for total monocytes, Ly6C-high and Ly6C-low monocytes at 24 hours after each treatment are shown. **B)** Fold change of monocyte subpopulations versus baseline after 6x R848 treatments. **C)** C57BL/6 mice or Ccr2RFP/RFP mice were either naive (WT n=4, Ccr2RFP/RFP n=4, grey bars) or received 6 R848 treatments (WT n=4, Ccr2RFP/RFP n=4, red bars). Immunofluorescent staining for CD68 (green) and nuclei (grey) was performed on tissue sections from lung and kidneys. Staining was quantified as mean CD68+ cells per field. Data representative of 2 (**C**) or 4 (**A-B**) independent experiments. Time course experiments analysed using two-way ANOVA with Tukey’s multiple comparisons test to compare between time points (**A**); for a single time-point one-way ANOVA with Bonferroni’s multiple comparison test for > 2 groups (**B, C**). Data are the mean ± SEM; only significant p values are indicated; * p<0.05 ** p<0.01, *** p<0.001, **** p<0.0001. Please also refer to Fig S5.

We next investigated CX3CR1, which has been shown to regulate peripheral levels of Ly6C-low monocytes (Landsman et al., 2009), using the *Cx3cr1^GFP/GFP^* knock-in mouse. At baseline, in the *Cx3cr1^GFP/GFP^* animals the peripheral Ly6C-low monocyte levels were approximately 50% of those in WT mice, as previously reported (Landsman et al., 2009). However, after 4x topical R848 treatments the *Cx3cr1^GFP/GFP^* mice developed a monocytosis that was skewed towards Ly6C-low subpopulation as in WT mice (**Fig S5A**) demonstrating that CX3CR1 is dispensable for R848-driven myelopoiesis. Given the independence of the monocyte egress from prototypical signals, we explored whether topical R848 could be promoting monocyte or haemopoietic progenitor mobilization in a non-targeted manner by increasing vascular permeability. We injected I.V into naïve and R848-treated mice the Evans Blue dye that binds to albumin and would leak into peripheral tissues during situations of decreased vascular integrity (Radu and Chernoff, 2013). We found that the Evans Blue amount in the ears from naïve mice was equal to that present in R848-treated ears and the contralateral ears of R848-treated mice (**Fig S5B**). In addition, there was no increase in bone marrow vascular permeability as assessed by Evans Blue quantity per tibia (**Fig S5C**). Thus, these findings demonstrate that the cutaneous TLR7 activation was able to stimulate the egression from the BM of monocytes circumventing the requirement of homeostatic cues like CCR2 or CX3CR1 by activating emergency pathways.

### The R848-induced emergency myeloid cells enhance viral control and limit the disease severity

As the BM egression of R848-induced monocytes occurred independently from the conventional regulatory mechanisms, we hypothesised these monocytes could be phenotypically and functionally distinct. Using the *Cx3cr1*^GFP/-^ mouse, Ly6C-low monocytes were visualized by intravital microscopy in the contralateral ear of mice treated topically 4x with R848. We observed a dramatic increase (∼10-fold) in intravascular CX3CR1-GFP+ monocytes in the R848-treated group (**Fig S6A**). These monocytes appeared to be rolling on the vessel wall indicating an activated phenotype (**Supp Video 1**). To confirm this, we comprehensively phenotyped the blood monocytes by flow cytometry. Consistent with the phenotype of the R848-induced bone marrow monocytes (**Fig 4E**), both blood subpopulations displayed expression of Sca1 (**Fig 7A**). In addition, both monocyte subpopulations expressed less F4/80 and more CD115 (**Fig 7B** and **Fig 7C**), suggesting an immature phenotype and a recent dependence on M-CSF. These changes were particularly pronounced in the Ly6C-low population, which appeared both activated and immature as reflected by the up-regulation of CXCR4, CD11c, CD11a and CD62L and the down-regulation of CCR2 (**Fig 7B**). Most notably, increased CXCR4 and decreased CCR2 expression phenocopy the changes seen in transitional BM pre-monocytes (Chong et al., 2016), which are not normally found in the blood, suggesting the premature egress of an emergency population. We next explored whether the R848-induced monocytes were also functionally impaired. We isolated BM monocytes from R848-treated and untreated mice and challenged them *in vitro* with R848 or LPS. We found that the R848-treated monocytes had a reduced cytokine response when rechallenged with the same TLR7 agonist but not with a different TLR stimulus (**Fig S6B**) suggesting a pathway-specific adaptation in the monocytes derived from the R848-exposed HSCs.

**Figure 7:**
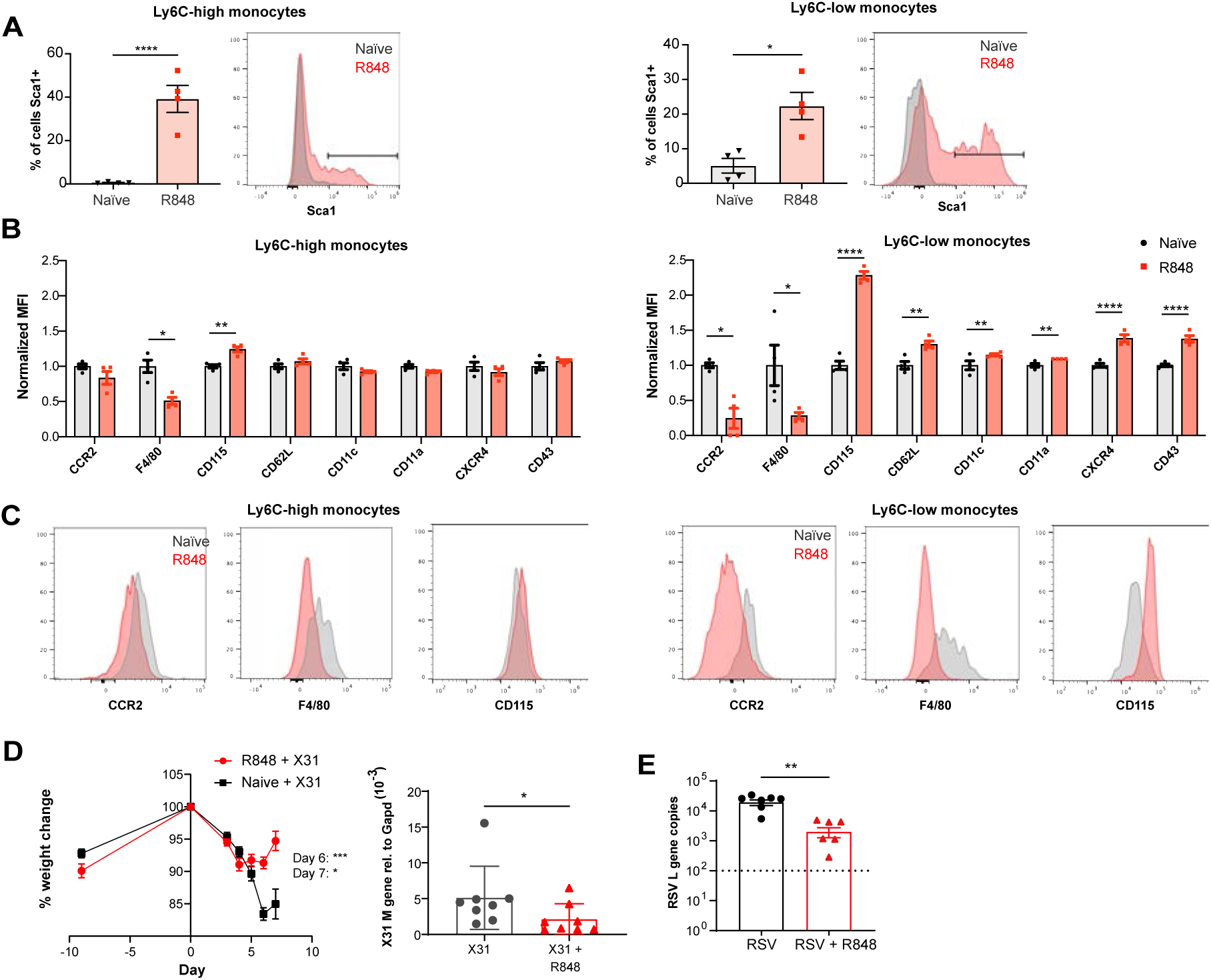
R848-induced emergency monocytes have anti-viral effects in the lung. **A-C)** C57BL/6 mice were either naive (n=4, black circles) or treated 4 times with topical R848 (n=4, red squares). **A)** Percentage of cells positive for Sca-1 and representative histograms in blood Ly6C-high and Ly6C-low monocytes from naive mice (grey) or R848-treated mice (red). **B)** Blood Ly6C-high and Ly6C-low monocytes were gated by flow cytometry and surface expression of the indicated proteins was quantified and expressed as MFI, normalized to the mean of the naive group. **C)** Representative histograms of CCR2, F4/80 and CD115 staining in blood Ly6C-high and Ly6C-low monocytes. **D)** Percentage of original weight (left panel) and lung viral load (right panel) in C57BL/6 mice (n=8/group) pre-treated 5 times with or without topical R848 infected intranasally with the influenza virus strains X31. Lung viral load at day 7 post inoculation was measured by quantification of matrix X31 gene copies in whole lung tissues. **E)** Lung viral load after RSV infection in C57BL/6 mice (n=6-7/group) pre-treated or not with topical R848 (5 times) was determined by quantification of viral L gene copies in lung tissues at day 4 post inoculation. Data representative of 2 independent experiments. Statistical analysis using unpaired t-test (**A-B; D right panel, E**); two-way ANOVA with Bonferroni’s post hoc test (**D, left panel**) Data are the mean ± SEM; only significant p values are indicated; * p<0.05, ** p<0.01, *** p<0.001, **** p<0.0001. Please also refer to Fig S6.

As the epithelial R848 applications promoted the migration and macrophage differentiation of Ly6C-high monocytes in peripheral organs like the lungs (**Fig 5C**), we next evaluated their contribution to control viral infections. We first infected R848-treated and control mice with the low pathogenicity H3N2 influenza strain X31 (Davidson et al., 2014). We found that the animals with the myeloid response showed reduced weight loss compared to untreated mice (**Fig 7D**). Consistent with the change in weight loss, the lung viral load, measured by expression of the influenza X31 matrix gene, was decreased in R848-treated mice (**Fig 7D**). Similarly, the extravasation of monocytes in the lung of R848-treated mice limited the viral replication after infection with respiratory syncytial virus (RSV) (**Fig 7E**), an RNA virus responsible for infant hospitalizations in the developed world.

Monocytes have previously been shown to have a beneficial anti-viral effect in the lungs (Goritzka et al., 2015). Together our findings indicate that the extravasation of monocytes into the lungs as result of the emergency monopoiesis triggered by cutaneous TLR7 activation indeed helps to limit viral replication and dampen disease severity. Altogether, this confirms the protective effect mediated by innate immunity even if the encounter with a pathogen-derived product occurs at a distant site.

## Discussion

Increased BM output of inflammatory cells, known as “emergency myelopoiesis”, is a critical feature of the host response to injury or infection. This process can be driven by systemic inflammatory factors and/or pathogen-derived products acting on precursor cells (Takizawa et al., 2012). Herein, we demonstrate that activation of TLR7 at the epithelial barrier of the skin and gut launches a specific and distinct monopoiesis response which does not occur after systemic TLR7 activation or activation of other TLRs. It is characterised by a rapid and dramatic increase of blood monocytes that egress from the BM bypassing the requirement of homeostatic cues like CCR2 or CX3CR1. The TLR7-induced monocytes appear to be immature and programmed to differentiate into macrophages, both surveying the endothelial integrity as Ly6C-low monocytes and trans-migrating into different organs to complement tissue resident macrophages. The response is orchestrated by BM-derived myeloid cells capable of transmitting the signals to the BM HSPCs independently from some of the well-known cytokines such type I and type II IFNs (summarised in the **Graphical Abstract**). Collectively our findings define the unique features of the BM myeloid response triggered by single-stranded RNA viruses on entering the epithelial barrier, a condition most likely occurring in infections with arboviruses and coronaviruses.

One of the key observations of our study is the unicity of the initial peripheral signal that is specific to, and shared between, epithelial barriers. Although the detailed mechanisms underlying this specificity remain to be defined, the canonical role of TLR7 in sensing viral ssRNA makes it evolutionarily logical for the response to be specific to epithelial barriers, which must be breached for infection to be established. The skin, lungs and the gut are similar in their constant exposure to mechanical trauma and a wide range of pathogens. To manage these insults, all these sites are equipped with an immune surveillance network comprising tissue resident macrophages, dendritic cells and mast cells; as well as resident lymphocytes such as γδ T cells and innate lymphoid cells. Notably, in the skin and gut a subset of macrophages are replenished by circulating monocytes (Bain et al., 2014; Kolter et al., 2019), unlike in other tissues where macrophages are embryonically derived, proliferate in situ and are only replenished by circulating monocytes after a severe insult (Ginhoux and Guilliams, 2016). Interestingly, TLR7 hyperactivation has been shown to drive a myeloid cell expansion in other settings, such as the transgenic over-expression of TLR7 in the TLR7.1 mouse (Deane et al., 2007). However, in these models the monocyte response appears to be driven by the extramedullary haematopoiesis in the spleen, whilst in our experimental model the BM is the only source of the myeloid expansion demonstrating the specificity of the peripheral cues induced by TLR7 activation in the skin and gut.

The monocytosis following topical R848 applications is accompanied by a dramatic expansion of HSPCs and monocyte progenitor cells in the BM and the egression in the peripheral blood of transitional BM pre-monocytes. Others have previously described a rapid expansion of phenotypically distinct Ly6C-high monocytes following infections with multiple intestinal pathogens, both parasitic and bacterial (Abidin et al., 2017; Askenase et al., 2015). In concordance with previous reports, we found that the expanded BM monocyte population expressed high levels of Sca-1 (Askenase et al., 2015). However, in our model the induced monocytosis was independent of both type I and type II IFNs indicating that a different priming mechanism had occurred. One interesting possibility is the type III IFN, which is increasingly recognised as an inducer of local anti-viral immunity at epithelial barriers (Lazear et al., 2015; Wack et al., 2015) and may have a still unrecognised role in the regulation of monocyte progenitors. It is also possible that CSF-1 signalling is involved in the monocyte expansion observed in our experimental model, given the up-regulation of the CSF1R (CD115) on the R848-expanded monocytes. Administration of recombinant CSF-1 can expand blood monocytes 5-10 fold after 10 days and requires repeated dosing (Ulich et al., 1990). However, it remains puzzling why CSF-1 secretion would only be initiated at epithelial barriers and not after systemic activation of TLR7 considering that this can induce extramedullary haematopoiesis in the spleen (Deane et al., 2007).

A striking feature of our model is the accelerated differentiation of Ly6C-high to Ly6C-low monocytes. Recent work has suggested that TLR7 activation of Ly6C-high monocytes using topical IMQ or *in vitro* can directly promote their conversion to Ly6C-low monocytes (Gamrekelashvili et al., 2020). However, while we agree that topical TLR7 activation can accelerate monocyte conversion, this appears to be an indirect effect driven by a secondary stimulus, as I.V or I.P administration of R848 did not result in an obvious monocyte ratio switch. As part of the enhanced differentiation program induced by topical R848, Ly6C-high monocytes also infiltrated peripheral organs and markedly increased the number of tissue macrophages with probably different consequences according to the organ. Global TLR activation using the TLR7.1 transgenic mouse has previously been shown to drive macrophage differentiation to inflammatory hemophagocytic macrophages which infiltrate the spleen and caused anaemia and thrombocytopenia (Akilesh et al., 2019). However, unlike the phenotype described here, this process was due to direct TLR7 activation in the BM and could be recapitulated using multiple daily I.P injection of R848, something that we did not detect using a less strong stimulation. In addition, in the paper by Akilesh *et al* (Akilesh et al., 2019) the generation of hemophagocytes could also be promoted by TLR9 activation, whereas we saw no effects of topical CpG application on peripheral monocytes. Whilst it is difficult to reconcile the different findings, one could argue that the TLR7.1 transgenic mouse and the hyper-stimulation applied by Akilesh *et al* in their model may recapitulate extreme phenotypes observed in the macrophage activation syndrome (MAS), a life-threatening complication of rheumatological diseases, whilst our experimental models mimic more closely the BM response to common single-stranded RNA viral infections.

The BM egression of monocytes in steady-steady conditions and during emergency myelopoiesis in response to inflammation and/or infections, including *Toxoplasma gondii* and *Listeria monocytogenes*, has been reported to depend on CCR2 expression (Grainger et al., 2013; Serbina and Pamer, 2006; Serbina et al., 2009) and this has become an accepted dogma in the literature. However, our findings demonstrate unequivocally that topical TLR7 application generates monocytes which are released from the BM in a CCR2-independent process. To our knowledge, the only other context in which this has been shown is during La Crosse virus infection, which interestingly has an ssRNA genome and is detected via TLR7 (Winkler et al., 2018). Consistent with a CCR2-independent egress, the R848-induced peripheral blood monocytes have low CCR2 expression and have an immature CXCR4-high, F4/80-low phenotype reminiscent of BM pre-monocytes (Chong et al., 2016). Under homeostatic conditions, CXCR4 and CCR2 play antagonistic roles dictating the respective retention or release of Ly6C-high monocytes from the BM (Jung et al., 2015). Therefore, the presence of up-regulated CXCR4 on blood monocytes potentially indicates their premature release to meet increased demand via a bypass of the CCR2/ CXCR4 axis. Of note, a potentially similar phenomenon has been suggested during SARS-CoV-2 infection, based on the presence of proliferation markers on blood monocytes (Mann et al., 2020). Given that SARS-CoV-2 has an ssRNA genome, further work is required to determine if our findings are applicable in this setting.

Another striking feature of our model was the accelerated extravasation of the immature R848-induced monocytes into multiple organs. Again, this egression occurred independently from conventional pathways like CCR2/CCL2 and was not associated with any obvious evidence of organ inflammation or damage, suggesting a distinct pathway. Consistent with the notion and previous data (Goritzka et al., 2015) that an early recruitment of Ly6C-high monocytes into the lung is a beneficial feature of the host innate immune response to virus infection, we found that the influx of R848-induced emergency myeloid cells restricted RSV and influenza virus replication and dampened disease severity. Similarly, monocyte-derived cells have been shown to limit HSV-2 and HSV-1 replication in the vaginal tract and in the cornea respectively (Conrady et al., 2013; Iijima et al., 2011), indicating that these cells play a key role in the resistance to viruses together with the type I IFNs. Notably, the fact that the monocytosis persisted long after the TLR7 stimulation at the epithelial barrier had ceased would suggest that the protective effects provided by these cells may prolong the antiviral effect of the type I IFNs. However, blood-derived tissue macrophages can also induce tissue damage. For example, inflammatory Ly6C-high monocytes have been shown to contribute to lung pathology in a model of influenza virus infection (Davidson et al., 2014; Herold et al., 2008; Lin et al., 2008), most likely recapitulating the pathology that occurs in severe infection if the initial innate response is dysregulated. A similar pattern has also been reported during SARS-CoV-2 infection where dynamic changes in the monocyte response have been correlated with COVID-19 disease severity (Schulte-Schrepping et al., 2020). Therefore, recruitment of antiviral monocytes during lung infections must be carefully balanced to reduce viral load but not allow excessive cell infiltration that can cause tissue damage.

In summary, the work presented here elucidates a novel pathway of emergency myelopoiesis which is uniquely activated by TLR7-expressing macrophages at epithelial barriers. Unlike in other settings, this is due to a specific expansion of the monocyte lineage from BM progenitors, is CCR2-independent and releases atypical pre-monocytes in the periphery. This emergency myeloid population is pushed towards a macrophage differentiation pathway which results in a predominance of Ly6C-low patrolling monocytes in the blood and a simultaneous infiltration of inflammatory macrophages into peripheral organs. Together these data challenge the dogma on how monocytes are produced during viral infections occurring at peripheral sites and highlight their contribution to the initial anti-viral immunity.

### Limitations of the study

While this study has revealed a novel pathway of emergency myelopoiesis uniquely triggered by TLR7 activation at epithelial barriers, it has a few limitations. First, we did not identify the mechanism(s) for the remote activation of monocytes. We assume that a soluble factor mediates this effect, but the nature of this soluble factor remains unclear. Additionally, whilst our study indicates that skin myeloid cells are responsible for this expansion of the monocyte lineage from BM progenitors, we could not pinpoint to a specific skin myeloid cell type. Furthermore, while our experiments demonstrate a CCR2-independent egression of BM immature monocytes as well as CCR2-independent migration of the Ly6C-high monocytes into different organs, we did not define the distinct pathways driving these processes. Moreover, how the R848-induced emergency myeloid cells modulate the lung anti-viral response remains a speculation and further studies will be needed to address these points.

## Material and Methods

### Mouse strains

All procedures were carried out in accordance with the institutional guidelines and the studies were approved by the UK Home Office. Experimental studies were designed according the ARRIVE guidelines (Percie du Sert et al., 2020). Experimental mice were 8-12 weeks of age, sex- and age-matched. All animals were housed in individually ventilated cages. The animals were selected randomly from a large pool, but no specific method of randomization was used to allocate mice into groups. The investigators were not blinded to allocation during experiment and outcome assessment.

BALB/c and C57BL/6 mice were purchased from Charles River (UK). The following mice were as previously described and maintained on a C57BL/6 background: B6.129P-*Cx3cr1*^tm1Litt/J^(*Cx3cr1^GFP/GFP^*) (Jung et al., 2000), *Ifnar1^-/-^* (Hwang et al., 1995), B6(Cg)-Rag2^tm1.1Cgn^/J (*Rag2^-/-^*) (Hao and Rajewsky, 2001), B6.129P2-Lyz2^tm1(cre)Ifo^/J (*Lyz2^Cre^*) (Clausen et al., 1999), B6.129(Cg)-Ccr2^tm2.1Ifc^/J (*Ccr2^RFP/RFP^*) (Saederup et al., 2010), B6.129P2-Tlr7^tm1Aki^ (*Tlr7^-/-^)* (Hemmi et al., 2002), B6.129S7-^Ifngtm1Ts^/J (*Ifng^-/-^*) (Dalton et al., 1993), *Tlr7^fl^* (Solmaz et al., 2019); HSC-SCL-Cre-ER^T^;R26R-EYFP (Göthert et al., 2005). The following mice were as previously described and maintained on a BALB/c background: *Gata1^tm6Sho^*/J (ΔdblGATA) (Yu et al., 2002), *Cpa3^Cre/+^* (Feyerabend et al., 2011).

### *In vivo* administration of TLR agonists

*Topical application*: mice were treated on the dorsal side of the right ear with 100 µg of R848 (Enzo Life Sciences, USA) diluted in 30 µl of acetone, 3 times per week up to a maximum of 12 treatments. The same dosing regimen was used for the following TLR ligands: Lipopolysaccharide (LPS) from E. coli 055:B5 (Invivogen), low molecular weight polyinosine-polycytidylic acid (Poly(I:C), Invivogen), Class C CpG oligonucleotide ODN-2395 (Invivogen). They were dissolved in 50:50 DMSO:PBS and applied topically at 100µg per dose. Imiquimod (IMQ) at a dose of 5 mg of 5% IMQ cream (Aldara^®^ cream─3M Pharmaceutical) was applied on the ventral side of one ear for 5 consecutive days. 5 nmol of 12-O-tetradecanoylphorbol-13-acetate (TPA) dissolved in ethanol was applied topically on the dorsal ear skin 4 times over 2 weeks.

*Other routes of administration:* i) mice treated intraperitoneally (I.P) received 100 µg of a water soluble R848 formulation (Invivogen) dissolved in 200 µl of PBS, 3 times per week; ii) mice treated intravenously received 100 µg of water-soluble R848 in 100µl of PBS, 3 times per week. Vehicle treated mice were used as controls; iii) mice treated orally received drinking water supplemented with R848 (Invivogen) at 6.3µg/ml to provide a calculated dose of ∼50µg/day/ mouse. Sodium saccharine sweetener (Sweetex) was added at 2 tablets per 100ml to improve palatability. Monocyte counts were monitored by flow cytometry of the peripheral blood at the indicated time points, as described in **Blood sampling.**

### Blocking and depletion experiments

*Blocking antibody*: mice were injected I.P with 10 mg of the anti-IL1R Anakinra (Kineret^®^, Sobi, Sweden) for 7 consecutive days, starting on the day of the first R848 treatment. Mice simultaneously received topical treatments every other day with either R848 or acetone. Mice were injected I.P with 200mg of the anti-TNF Etanercept (Enbrel^®^) or PBS for 9 consecutive days, starting two days prior to the first R848 treatment. Mice were injected I.P with 200 mg of the anti-IL-6R (Clone 15A7, BioXcell) or isotype control (BioXcell) or PBS every three days, starting one day prior to the first R848 treatment. Mice then received topical R848 treatments every other day for 4 times . Monocyte counts were assessed by flow cytometry at the indicated time points, as described in **Blood sampling** *Neutrophil depletion*: mice were injected I.P with 200µg of a chimeric anti-Ly6G antibody composed of the 1A8 variable region with a mouse IgG2a Fc region, to avoid eliciting a neutralizing response (Absolute Antibody) (Daley et al., 2008). Mice received 4 topical R848 treatments at a frequency of 3 treatments per week and antibody was injected 4 hours prior to each R848 treatment. Blood neutrophil depletion was validated by flow cytometry, with neutrophils defined independently of their Ly6G expression as SSC^high^, CD11b^pos^, Ly6C^int^, CD115^neg^.

*Ly6C-high monocyte depletion:* mice were injected I.P with 100µg of a chimeric anti-GR1 antibody composed of the RB6-8C5 variable region with a mouse IgG2a Fc region, to avoid eliciting a neutralizing response (Absolute Antibody). Mice received 4 topical R848 treatments at a frequency of 3 treatments per week and antibody was injected on 3 consecutive days, starting with the 3^rd^ R848 treatment. Blood monocyte depletion was validated by flow cytometry at 24 hours after each antibody injection.

### Splenectomy

Eight week-old BALB/c mice underwent surgical splenectomy or sham surgery under aseptic conditions. Briefly, mice were injected subcutaneously with ketamine (80 mg/kg) and xylazine (16 mg/kg) before being anaesthetised with 5% Isofluorane. Mice were rested for 6 weeks before being treated 6 times with 100µg topical R848 over 2 weeks. To assess monocytosis, a blood sample was taken prior to the first R848 treatment and at 24 hours after each treatment.

### Intravital microscopy

*Cx3cr1* ^GFP/+^ mice, 10-12 weeks of age, were treated topically for 4 times with R848 or acetone on one ear, before being used for imaging. Intravital imaging of the dermis of the contralateral ear was performed as previously described (Carlin et al., 2013). Briefly, after I.P administration of anaesthetic cocktail of fentanyl/fluanisone and midazolam, mice were maintained at 37°C with oxygen supplementation. 80μL of tetramethylrhodamine (TRITC) conjugated 70kDa dextran (70μM) was injected I.V and the ear to be imaged was taped to the centre of the coverslip. Light was generated from 488-nm and 562-nm lasers, emitted light signal was detected to generate two colour 8-bit images using a 10x/0.4 objective on a Leica SP5 confocal microscope. Images were analysed using Imaris software (Bitplane). Dextran signal was used to identify the intravascular cells and monocytes were automatically selected on the base of the quality and intensity of their GFP signal.

### BrDU incorporation

C57BL/6 mice were treated twice with topical R848 and then were injected once I.P with 2 mg of (5-bromo-2’-deoxyuridine) BrdU (Biolegend). Bone-marrow (BM) was harvested 16 hours later and samples were processed with BrdU staining kit (e-Bioscience, USA) according to the manufacturer’s provided protocol and analysed by flow-cytometry (see details in Methods).

### Lineage tracing experiment

HSC-SCL-Cre-ER^T^;R26R-EYFP mice were given 100μL tamoxifen (40mg/mL) by oral gavage for 5 consecutive days. After 3 days, some mice received topical R848 treatments every other day for 4 times and others were left untreated. Monocyte counts were assessed by flow cytometry at the indicated time points, as described in **Blood sampling**.

### Bone Marrow Chimeras

Prior to irradiation, all mice were treated orally with Baytril (Bayer) for 7 days and moved to sterile cages. C57BL/6 or *Tlr7^-/-^* host mice were lethally irradiated (700cGy) and reconstituted I.V on the same day with 5 x 10^6^ BM cells from either C57BL/6 or *Tlr7^-/-^* donor mice. Mice were left to recover for 8 weeks and reconstitution was confirmed by blood sampling and flow cytometry. Reconstituted mice were treated topically with 100µg R848, 3 times per week for 4 treatments.

### Vascular Permeability

BALB/c mice were naive or treated topically 4 times with 100µg R848. 200µl of 0.5% Evans Blue in PBS was injected I.V at 24 hours after the final R848 treatment. Mice were culled by cervical dislocation after 30 minutes and ears and right tibia were harvested. Ears were minced into a slurry and tibias were crushed, before adding each to 500µl formamide. After 48 hours of incubation at room temperature, the absorbance of the formamide was read at 620nm and concentration of Evans Blue was calculated using a standard curve.

### Virus infections and qPCR

C57BL/6 were naive or treated topically 5 times with 100µg R848 prior to infection with the influenza A virus strain X31 (obtained from John McCauley, The Francis Crick Institute, UK) or plaque-purified human RSV (originally A2 strain from ATCC, USA, grown in HEp2 cells). The topical application of R848 was continued during the viral infection. For infections mice were lightly anaesthetized (3% Isofluorane) and virus was intranasally (I.N) administered. Mice received 8×10^5^ focus forming units (FFU) of RSV or 250 plaque forming units (PFU) of X31, both given in 100 µl. Weight was monitored daily. Mice were sacrificed at day 7 (X31) and day 4 (RSV) post infection (p.i.) and lung lobes were stored in RNA-later (Sigma). Lung tissue was homogenized using a TissueLyser LT (Qiagen), and total RNA was extracted using RNeasy Mini kit including DNA removal (Qiagen) according to the manufacturer’s instructions. 1-2 µg of RNA was reverse-transcribed using the High-Capacity RNA-to-cDNA kit according to the manufacturer’s instructions (Applied Biosystems). qPCR was performed to quantify lung RNA levels using the mastermix QuantiTech Probe PCR kit (Qiagen). To quantify RSV L genes, primers and FAM-TAMRA probes previously described were used (Lee et al., 2010). The absolute number of gene copies was calculated using a plasmid DNA standard curve and the results were normalized to levels of *Gapdh* (Applied Biosystems). The relative quantification of X31 M genes (primers from Ward et al., 2004) was expressed relatively to the expression of *Gapdh*. First, the ΔCt (Ct = cycle threshold) between the target gene and the *Gapdh* for each sample was calculated, then the expression was calculated as 2^-ΔCt^. Analysis was performed using 7500 Fast System SDS software (Applied Biosystems).

### *In vitro* stimulation

C57BL/6 mice were naïve or treated topically 4 times with 100µg R848. Femur and tibia were harvested without breaking the bone, to maintain sterility. Collected bones were placed into RPMI 1640 (Gibco), 10% FBS, 100 U/mL penicillin, and 100 g/mL streptomycin. Bones were cleaned and flushed with HBSS-2% FCS and cells were pooled from 2 mice per biological replicate. Cell suspensions were washed and re-suspended in hypotonic red blood cell lysis buffer (RBC Lysis Buffer: 1L of water, 8.02g NH4Cl, 0.84g NaHCO3, 0.37g EDTA) for 5 minutes on ice. Cells were wash and re-suspended in PBS solution containing 1% BSA, 2mM EDTA (Sorting buffer) and cell surface staining was then performed using the fluorochrome-conjugated antibodies as in **Flow Cytometry**. Monocytes were sorted into HBSS-2% FCS in 15ml Falcon tubes using a BD Aria III (BD Biosciences). Monocytes were gated as Lineage-negative (B220, CD49b, Ly6G), CD115-positive, CD117-negative and CD11b-positive. Sorted cells were washed and re-suspended in DMEM (Gibco), 20% FCS, 100 U/mL penicillin, and 100 g/mL streptomycin, 2mM L-glutamine at 1×10^6^ per ml. 1×10^5^ cells per replicate were stimulated with either 100ng/ml R848 (Invivogen) or 1µg/ml LPS (O127:B8, Sigma) or medium alone for 16 hours. Supernatant was harvested and cytokines were quantified using a Legendplex Mix and Match Kit according to manufacturer instructions (Biolegend).

## METHOD DETAILS

### Flow cytometry

Where Live/Dead staining was performed, single cell suspensions were stained for 20 minutes at room temperature with Live/Dead Fixable Dead Cell Stain (Life), at the dilution of 1/1000 in PBS, as per manufacturer protocol. Cells were incubated with a saturating concentration of 2.4G2 monoclonal antibody (anti-CD16/32) to block non-specific Fc receptor binding and stained in PBS solution containing 1% BSA, 2mM EDTA, and 0.09% NaN3 (FACS buffer) with an appropriate dilution of fluorophore-conjugated antibodies for 20 minutes at 4°C (see antibodies used in **Key Resources Table**). Samples were acquired on either a Fortessa X20 (BD Biosciences) or an Aurora (Cytek) flow cytometer and analysed using FlowJo X for Mac (TreeStar).

### Blood sampling

Tail vein blood samples were added to an equal volume of 100mM EDTA, washed and re-suspended in FACS buffer. Cell surface staining was then performed using the fluorochrome-conjugated antibodies listed in **Flow Cytometry**. After antibody staining, cells were washed and red blood cells were removed by hypotonic lysis using BD FACS Lysing Solution (BD Biosciences). Absolute cell counts were quantified with AccuCheck counting beads (ThermoFisher Scientific, USA) as per manufacturer’s protocol. Total blood monocytes were gated as follows and as illustrated in **Figure 1C**: CD3-neg, B220-neg, CD11b+, CD115+. Monocytes were then split into Ly6C-high and Ly6C-low subpopulations based on their expression of Ly6C and CD11a.

### Bone marrow staining

Femur and tibia were harvested without breaking the bone, to avoid blood contamination. Collected bones were placed into RPMI 1640 (Gibco), 10% FBS, 100 U/mL penicillin, and 100 g/mL streptomycin. Bones were cleaned and flushed with HBSS-2% FCS, and cell suspensions were washed and re-suspended in hypotonic BD FACS Lysing Solution (BD Biosciences) for 2 minutes on ice. Cells were washed and resuspended in FACS buffer. Antibody staining was performed as described above in **Flow cytometry.** The gating for BM populations was adapted from a published strategy (Yáñez et al., 2017) and is illustrated in **Supp** **Fig 4A**. All populations are first gated lineage-negative (CD3, B220, Ly6G, CD49b). Haematopoietic stem and progenitor cells, HSPC: CD115-neg, c-kit+, Sca-1-neg; monocyte-dendritic cell progenitor, MDP: CD115+, c-Kit+, Ly6C-low; common monocyte progenitor, <u>cMoP:</u> CD115+, c-Kit+, Ly6C-high, Flt3-neg; granulocyte-monocyte progenitor, GMP: CD115-neg, cKit+, Sca-1-neg, CD16/ 32+, Ly6C-low; common myeloid progenitor, CMP: CD115-neg, cKit+, Sca-1-neg, CD16/ 32-neg, Ly6C-low; granulocyte-committed progenitor, GP: CD115-neg, cKit+, Sca-1-neg, CD16/ 32+, Ly6C-high; Ly6C-high monocytes: CD115+, c-Kit-neg, Ly6C-high; Ly6C-low monocytes: CD115+, c-Kit-neg, Ly6C-low.

### Skin digestion

Single cell suspensions from ear skin was obtained as follows: i) ears were split dorsal-ventral using forceps to analyse only the dorsal (treated) side; ii) skin was cut into ∼1mm^2^ pieces using a scalpel blade and incubated for 2 hours in digestion buffer containing 25 μg/ml Liberase (Roche), 250 μg/ml DNAseI (Roche) and 1 × DNAse buffer (1.21 Tris base, 0.5 g MgCl2 and 0.073 g CaCl2) at 37°C; iii) pieces were transferred into C-tubes (Miltenyi Biotech) containing RPMI-1640 medium (Thermo Fisher) supplemented with 10% heat-inactivated FCS and physically disrupted using a GentleMACS dissociator (Miltenyi); iv) cell suspensions were then filtered through 70µM cell strainers (BD Bioscience) and counted using a CASY cell counter (Roche). Cell suspensions were stained as described above in **Flow Cytometry**.

### Immunohistochemistry

Lungs, livers and kidneys were harvested and snap frozen in OCT using isopentane cooled to -80°C on dry ice. 9µM sections were cut using a Leica JUNG CM1800 cryostat and stored at – 80°C. For staining, slides were equilibrated to room temperature before fixation in ice-cold acetone for 10 minutes. Sections were blocked in PBS with 5% BSA and 10% serum (dependent on the species of the secondary antibody being used) for 1 hour at room temperature. All antibody staining steps were performed in PBS-5% BSA-0.1% Triton X-100 for 45 minutes at room temperature. The following primary antibodies were used: anti-CD68 AlexaFluor 488 (FA-11, Biolegend), anti-F4/80 AlexaFluor 488 (BM8), anti-CD43 (S7, BD Biosciences). Hoescht 33342 (NucBlue, ThermoFisher) was added to the final staining step according to manufacturer instructions and sections were mounted in ProLong Glass (ThermoFisher). Images were acquired on a Zeiss Axio Observer inverted widefield microscope using a Colibri.2 LED illumination source, a 20X/ 0.8 plan-apochromat objective and a Hamamatsu Flash 4.0 camera. Images were processed to correct brightness and contrast in FIJI (Schindelin et al., 2012). Cell quantification was performed using the surfaces function of Imaris 8 (Bitplane), with a minimum of 5 randomly selected fields per section being used for analysis. Data are expressed as the mean number of cells per field.

## QUANTIFICATION AND STATISTICAL ANALYSIS

Statistical comparisons between two groups at a single time point were performed using a two-tailed, unpaired t-test. In experiments with more than two groups at a single time point, analysis was by one-way ANOVA with Tukey’s multiple comparisons test. For data sets with multiple groups over a time-course, analysis was performed using two-way ANOVA with either Tukey’s or Bonferonni’s multiple comparisons test as appropriate and indicated in the figure legends. Statistical analysis was performed using Prism 9.0 (GraphPad).

## Acknowledgments

We thank Tim Sparwasser for providing the *Tlr7*-floxed mice. We are indebted to the staff of the Imperial Central Biomedical Services for the care of the animals and the LMS/NIHR Imperial Biomedical Research Centre Flow Cytometry Facility for FACS support. CG was supported by an award from Imperial Institutional Strategic Support Fund. The research reported herein was supported by the Wellcome Trust (Grant reference number: 108008/Z/15/Z (to M.B.) and 102126/B/13/Z (to CJ:AO). CG was supported by an Imperial Wellcome Trust Institutional Strategic Support Fund and TCL by a Sir Henry Dale Fellowship from the Wellcome Trust and Royal Society (210424/Z/18/Z) and a KKLF project grant (KKL1379). We acknowledge contribution from the National Institute for Health Research (NIHR) Biomedical Research Centre based at Imperial College Healthcare NHS Trust and Imperial College London. The views expressed are those of the author(s) and not necessarily those of the NHS, the NIHR or the Department of Health.

## Author contributions

Conceptualization: WDJ, CG, JS, KJW, MB.; Methodology: WDJ, CG, TCL.; Investigation: WDJ, CG, SW, AO, SS, TCL.; Resources: TCL, CJ.; Writing-Original Draft: WDJ, MB.; Writing-Review & Editing: WDJ, CG, KJW, JS, MB.; Visualization: WDJ, CG; Supervision: CJ, JS, KJW, MB.; Funding Acquisition: MB.

---

**Supplemental Figure 1:**
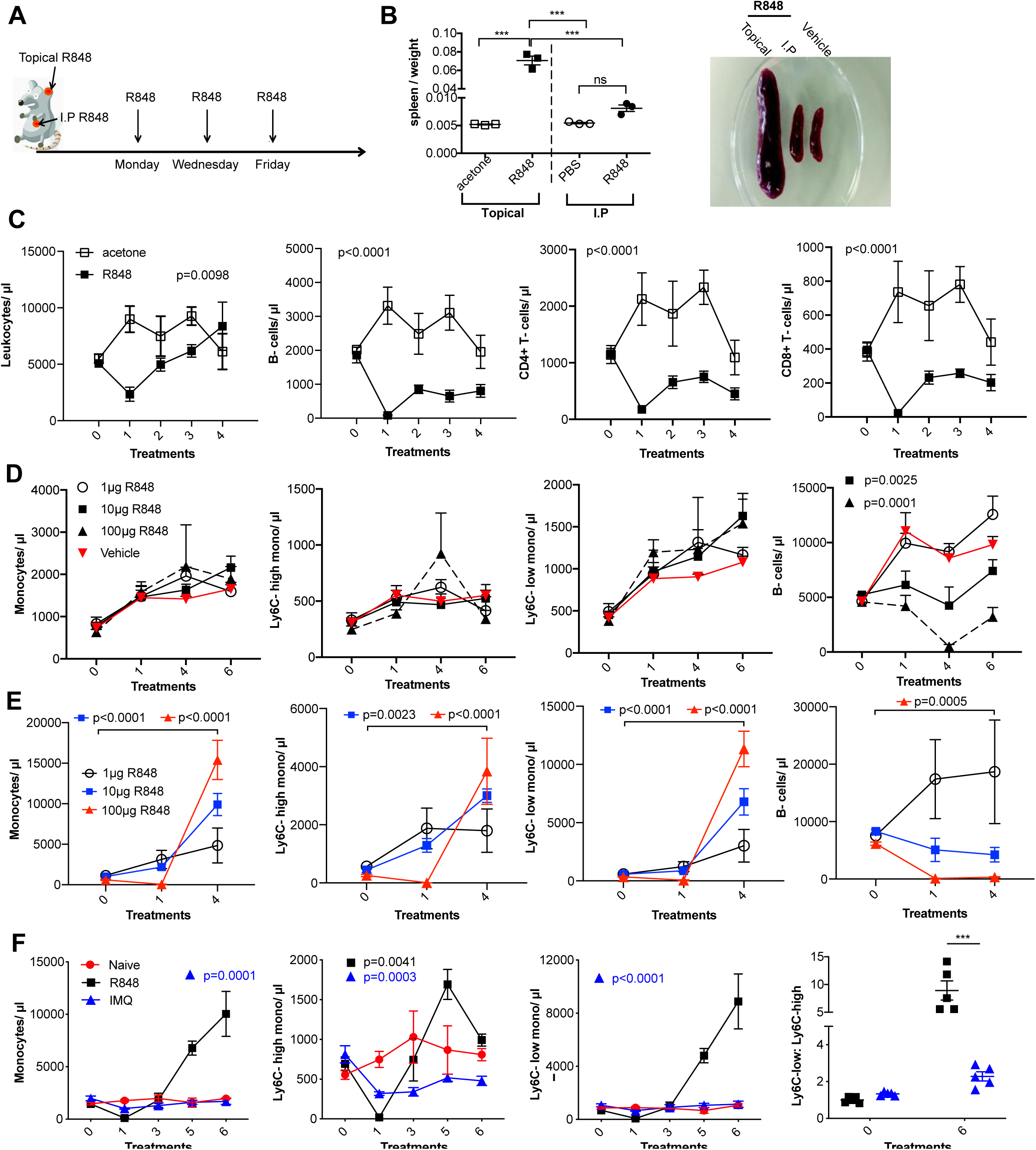
The effects of TLR7 stimulation depend on dose and route of administration. **A)** Graphical illustration of R848 treatment regimen. **B)** BALB/c mice (n=3 per group) received 100μg of R848 topically or I.P, 3x per week for 4 weeks. Control mice given topical acetone or 200μl of PBS I.P. Splenomegaly quantified as spleen: body weight ratio, with representative example shown. **C)** BALB/c mice (n=3 per group) received 4 treatments with 100μg topical R848 (black squares) or acetone (open squares). Blood counts are shown (left to right) for total leukocytes, B-cells, CD4+ T-cells and CD8+ T-cells 24 hours after each treatment. **D-E**) Blood counts for total monocytes, Ly6C-high monocytes, Ly6C-low monocytes and B-cells 24 hours after each treatment. **D)** C57BL/6 mice (n=4 per group) received 6x I.P treatments with 1μg (open circles), 10μg (black squares) or 100μg R848 (black triangles) or PBS vehicle (red triangles). Statistics versus vehicle group. **E)** C57BL/6 mice (n=4 per group) received 4x topical treatments with 1μ g (open circles), 10μg (blue squares) or 100μg R848 (red triangles). Statistics show pre-bleed vs 4 treatments. **F)** C57BL/6 mice were either naive (red circles, n=3) or received 6 daily topical treatments with imiquimod (IMQ, blue triangles, n=5) or R848 (black squares, n=5). Blood counts 24 hours after the indicated treatments. Statistics versus naive group. Data representative of 2 experiments (except **B** and **E**). One-way ANOVA with Bonferroni’s multiple comparison test for a single time point (**B**); two-way ANOVA with Tukey’s multiple comparisons test for analysis of time-course experiments (**C-F**). Data are the mean ± SEM; only significant p values are indicated; ***p<0.001.

**Supplemental Figure 2:**
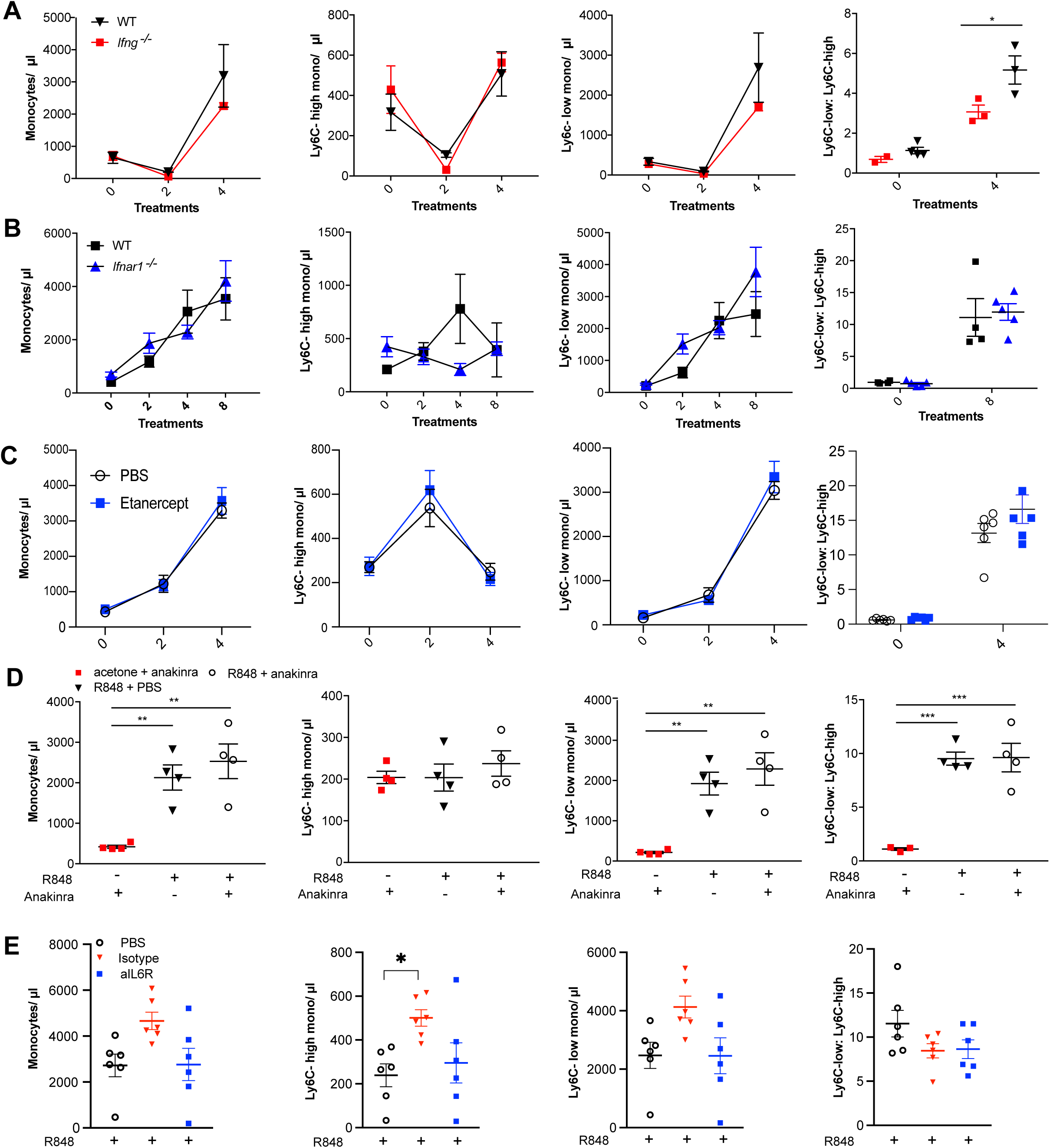
IFNs and cytokines are not involved in the R848-induced monocytosis. **A)** C57BL/6 (black triangles) or Ifng-/-mice (red squares) were either left untreated (n=4 WT, n=2 Ifng-/-) or given 4x topical treatments with R848 (n=3 per group). Blood counts for total monocytes, Ly6C-high monocytes, Ly6C-low monocytes and the monocyte subpopulation ratio at baseline and at 24 hours after the indicated treatment. Data representative of 2 independent experiments. **B)** C57BL/6 (n=4, black squares) or Ifnar1-/-(n=5, blue triangles) mice received 8 topical treatments with R848. Blood counts for total monocytes, Ly6C-high monocytes, Ly6C-low monocytes and the monocyte subpopulation ratio at baseline and 24 hours after the indicated treatments. **C)** BALB/c mice were treated daily I.P with Etanercept (anti-TNFalfa) (n=6, blue squares) or PBS (n=6, open circles) and then received 4x topical R848 treatments. Blood counts for total monocytes, Ly6C-high monocytes, Ly6C-low monocytes and the monocyte subpopulation ratio at baseline and at 24 hours after the indicated treatment. **D)** BALB/c mice received the following combinations: topical R848 and anakinra daily I.P (n=4, open circles), topical R848 and daily PBS I.P (n=4, black triangles), topical acetone and anakinra daily I.P (n=4, red squares). Blood counts for total monocytes, Ly6C-high monocytes, Ly6C-low mono-cytes and the monocyte subpopulation ratio at baseline and at 24 hours after the 4th treatment of R848. **E)** BALB/c mice were treated I.P with PBS (n=6, open circles) or anti-IL6R (n=6, blue squares) or the isotype control (n=6, red triangle) every 3 days and received 4x topical R848 applications. Blood counts for total monocytes, Ly6C-high monocytes, Ly6C-low monocytes and the monocyte subpopulation ratio at baseline and at 24 hours after the 4th treatment with R848. Time course experiments analysed using two-way ANOVA with Tukey’s multiple comparisons test to compare between time points (A-C); one-way ANOVA with Bonferroni’s multiple comparison test for comparing >2 groups at a single time-point calculated (D, E). Data are the mean ± SEM; only significant p values are indicated; * p<0.05, ** p<0.01, *** p<0.001.

**Supplemental Figure 3:**
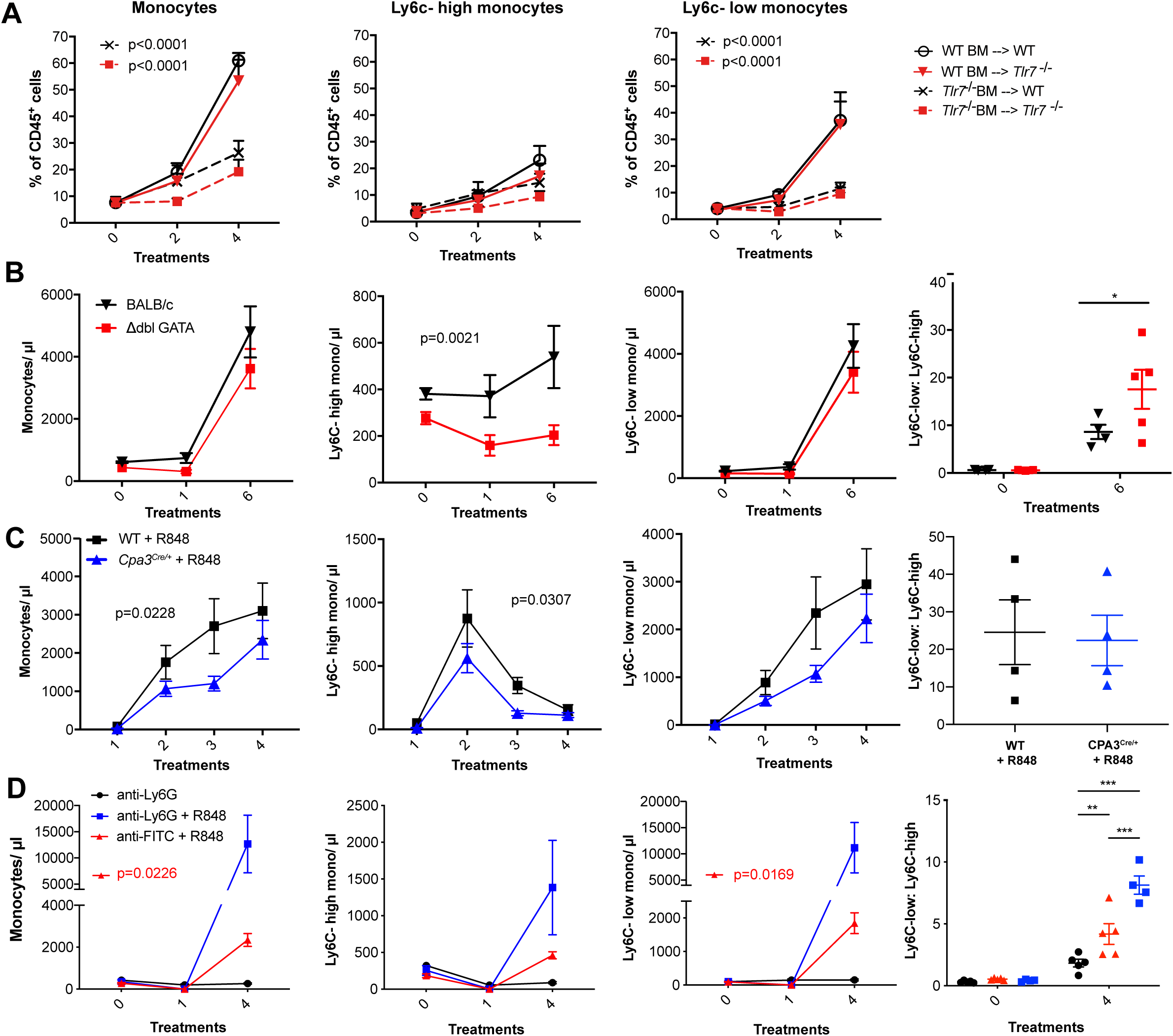
The role of myeloid cell populations in R848-driven monocytosis. **A)** Lethally irradiated WT or Tlr7-/-host mice were reconstituted with BM from WT or Tlr7-/-donor mice as follows: WT BM into WT host (n=5, open circles), WT BM into Tlr7-/-host (n=5, red triangles), Tlr7-/-BM into WT host (n=5, crosses), Tlr7-/-BM into Tlr7-/-host (n=5, red squares). Each group received 4x topical treatments with R848. Total blood monocytes, Ly6C-high monocytes and Ly6C-low monocytes are shown as a proportion of CD45+ cells, at baseline and 24 hours after the indicated treatments. **B-D)** Blood counts for total monocytes, Ly6C-high monocytes, Ly6C-low monocytes and the monocyte subpopulation ratio at baseline and 24 hours after the indicated R848 treatments. **B)** BALB/c (n=4, black triangles) or ΔdblGATA (n=6, red squares) mice received 6 topical treatments with R848. **C)** BALB/ c mice were either treated with topical R848 (n=4, black squares) alongside a group of Cpa3-Cre-/+ mice (n=4, blue triangles). **D)** BALB/c mice were treated with combinations of topical R848 and I.P anti-Ly6G or anti-FITC isotype control, with the antibody injected 4 hours prior to the first R848 treatment. Treatments were in the following groups: isotype control and R848 (n=5, red triangles), anti-Ly6G and R848 (n=4, blue squares), anti-Ly6G alone (n=5, black circles). Statistics compare anti-Ly6G+R848 to anti-FITC+R848. Time-course experiments analysed using two-way ANOVA with Tukey’s multiple comparisons test to compare between groups at a given time-point (**A-D**); one-way ANOVA with Bonferroni’s multiple comparison test for comparing >2 groups at a single time-point; unpaired t-test for comparing two groups at a single time point. Data are the mean ± SEM; only significant p values are indicated;* p<0.05, ** p<0.01, *** p<0.001.

**Supplemental Figure 4:**
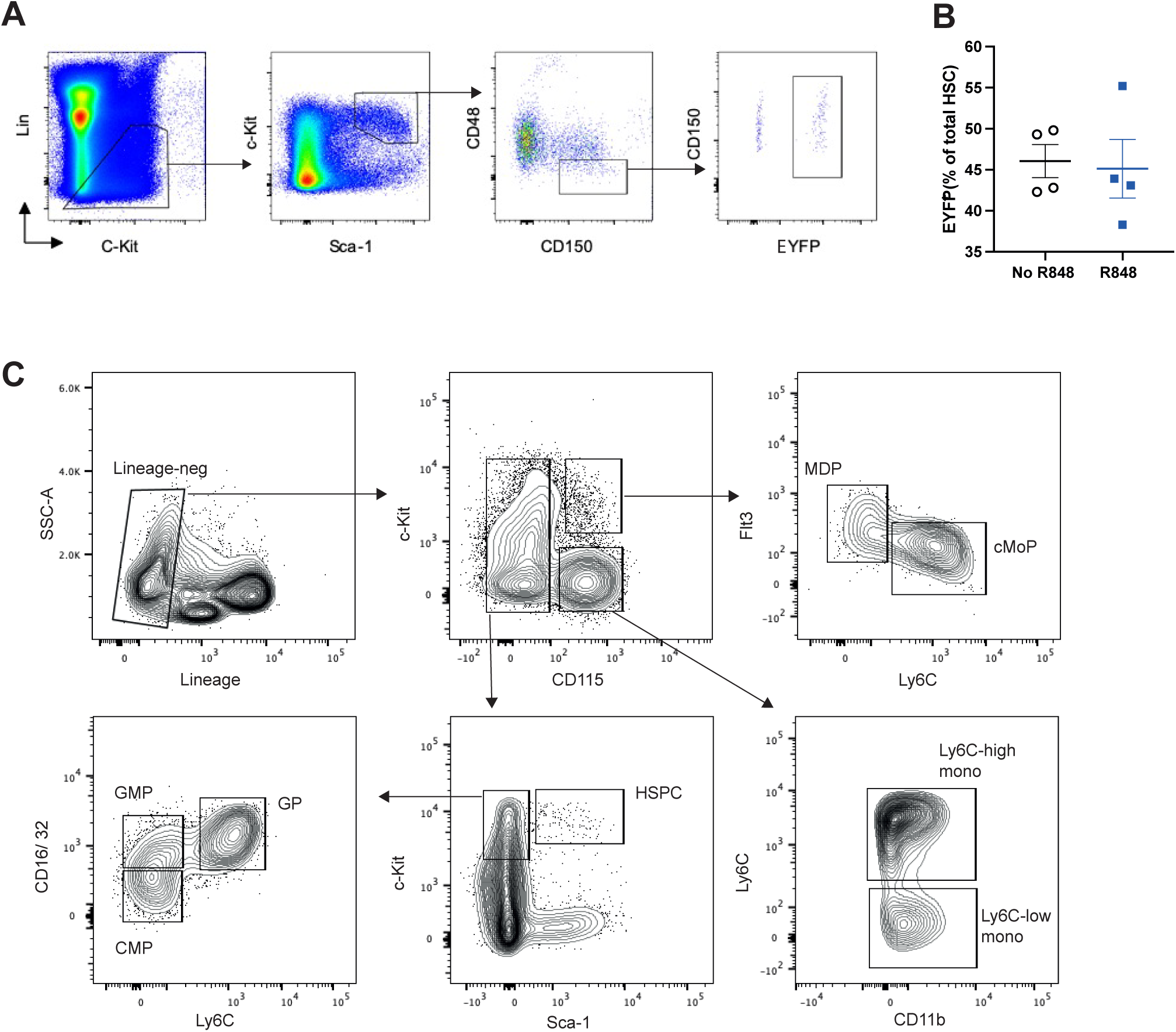
Bone marrow gating strategy. A-B) HSC-SCL-Cre-ERT;R26R-EYFP mice (n=4 per group) were treated for 5 consecutive days with Tamoxifen (4mg/100μL) by oral gavage. Three days after the tamoxifen administration, the left ears were treated topically 4x with R848 or left untreated. A) Gating strategy for YFP+HSCs in bone marrow. HSCs are excluded for doublets using side scatter area and side scatter height prior and gated lineage negative (B220, CD5, CD4, CD8, TER119 and GR-1). HSCs were then defined as c-Kit+ Sca-1+ CD150+CD48-. EYFP was then gated to determine recombination efficiency. B) Percentage of EYFP+HSCs within the HSCs in the BM after tamoxifen and 4xR848 treatment (R848 group) or tamoxifen only (no R848 group). Representative of 2 independent experiments. C) Gating strategy for BM analysis. All populations are gated lineage-negative (CD3, B220, Ly6G, CD49b). HSPC: CD115-neg, c-kit+, Sca-1-neg; MDP: CD115+, c-Kit+, Ly6C-low; cMoP: CD115+, c-Kit+, Ly6C-high, Flt3-neg; GMP: CD115-neg, cKit+, Sca-1-neg, CD16/ 32+, Ly6C-low; CMP: CD115-neg, c-Kit+, Sca-1-neg, CD16/ 32-neg, Ly6C-low; GP: CD115-neg, cKit+, Sca-1-neg, CD16/ 32+, Ly6C-high; Ly6C-high monocytes: CD115+, c-Kit-neg, Ly6C-high; Ly6C-low monocytes: CD115+, c-Kit-neg, Ly6C-low. Haematopoetic stem and progenitor cells: HSPC; common myeloid progenitor: CMP; monocyte-dendritic precursor: MDP; granulocyte-monocyte progenitor: GMP; common monocyte progenitor: cMoP; common dendritic progenitor: CDP: monocyte-committed progenitor: MP; granulocyte-committed progenitor: GP

**Supplemental Figure 5:**
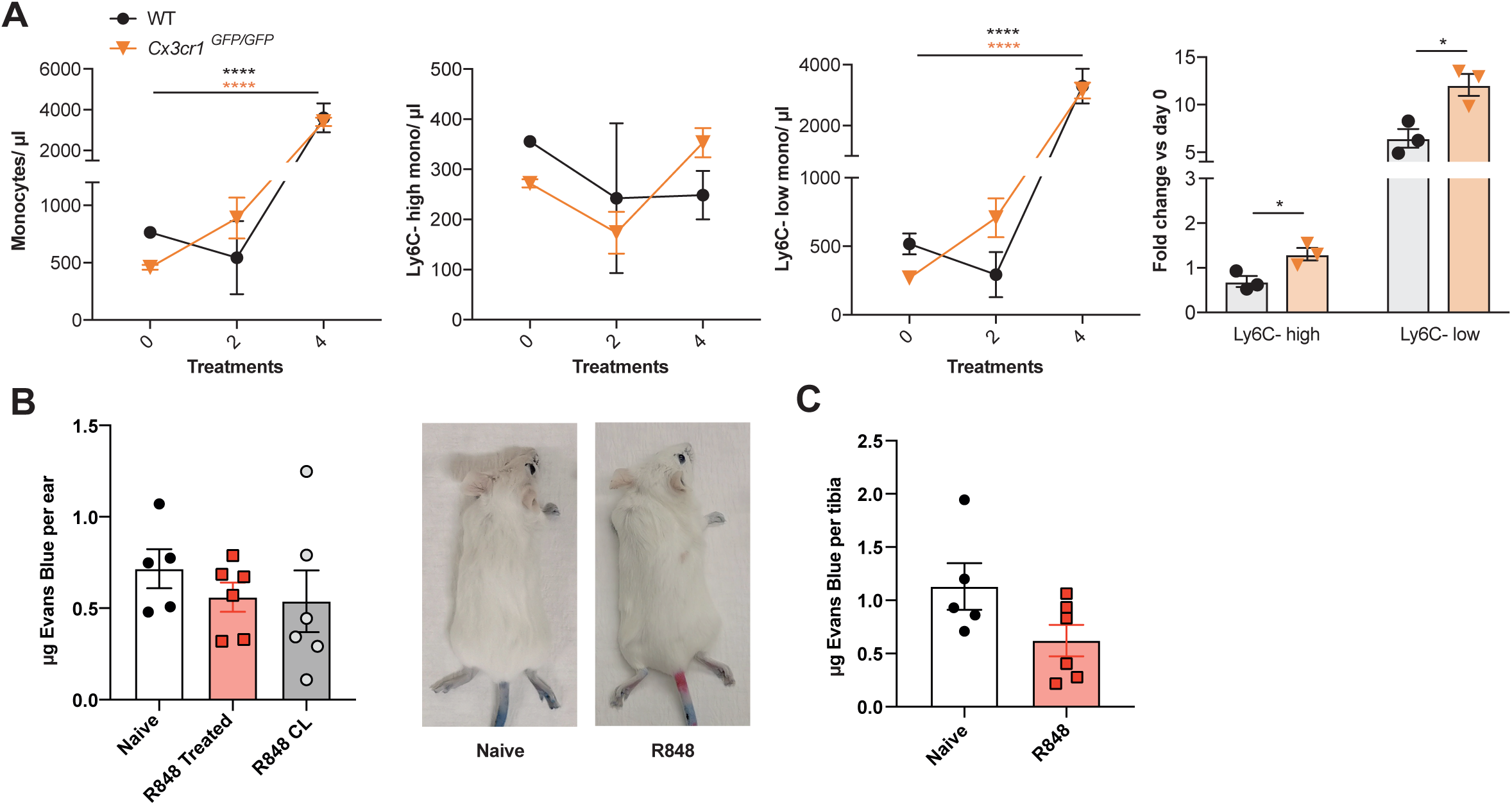
R848-induced myeloid response is independent from CX3CR1 and vascular permeability. **A)** C57BL/6 mice (n=3, black lines) or Cx3cr1GFP/GFP (n=3, orange lines) were treated topically 4 times with R848. Blood counts for total monocytes, Ly6C-high monocytes, Ly6C-low monocytes and the fold change of monocyte subpopulations after 4 treatments versus baseline. **B-C)** BALB/c mice were naive (n=5) or treated topically 4 times with R848 (n=6). 1mg Evans Blue was injected I.V at 24 hours after the final R848 treatment. **B)** Quantity of Evans Blue per ear is shown for naive mice (white bars), the R848-treated ear (red bars) and the contralateral ear (grey bars), along with representative photographs. **C)** Evans Blue is quantified as µg per tibia in naïve and R848-treated mice. Data representative of 2 independent experiments. Time course experiments analysed using two-way ANOVA with Tukey’s multiple comparisons test to compare between time points (A); fold change data analysed using unpaired t-test (**A**); two groups at a single time point compared using unpaired t-test (**C**); one-way ANOVA with Bonferroni’s multiple comparison test for >2 groups at a single time-point (**B**). Data are the mean ± SEM; only significant p values are indicated; * p<0.05, **** p<0.0001.

**Supplemental Figure 6:**
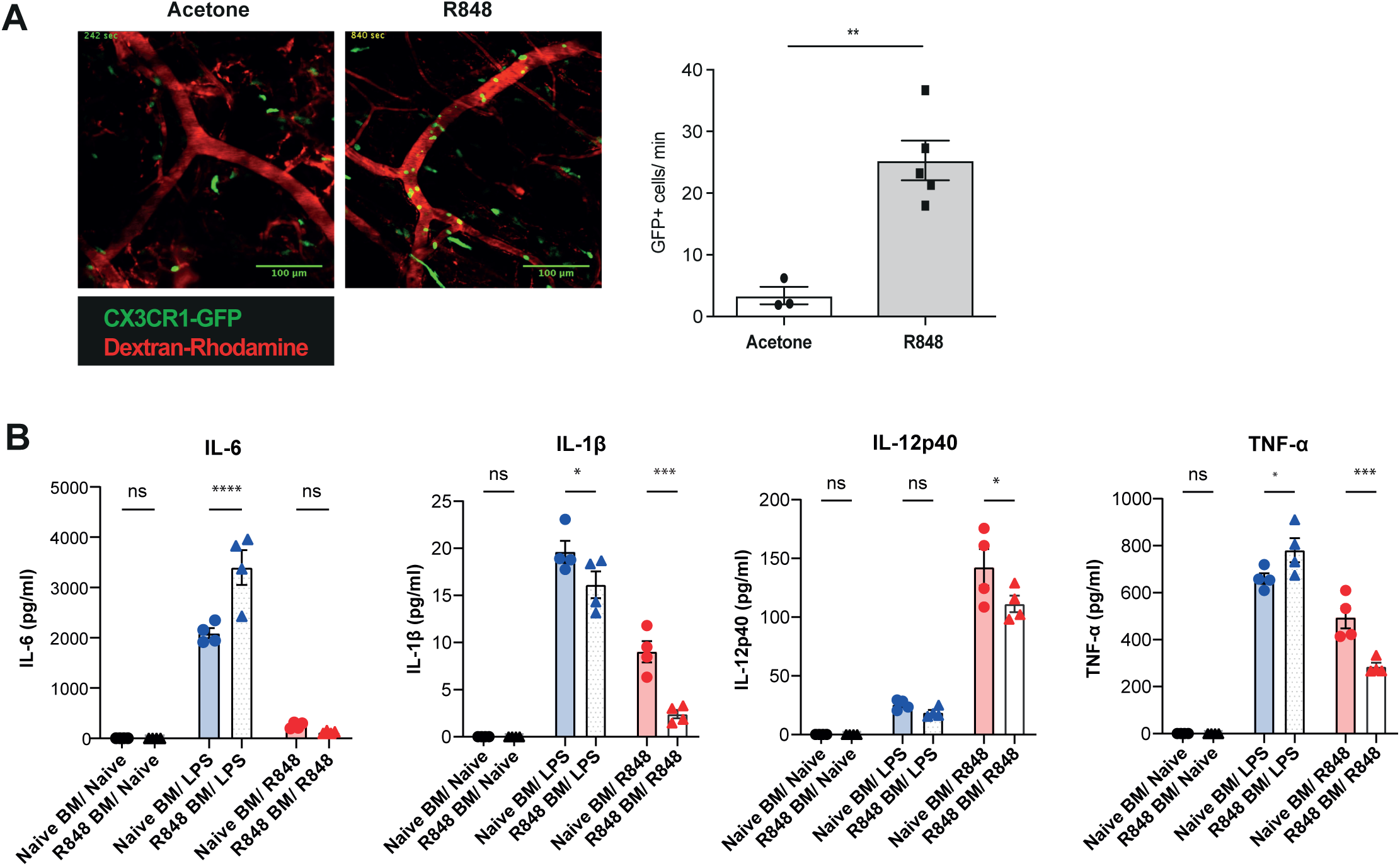
R848-induced monocytes are activated and functionally impaired. A) Cx3cr1GFP/+ mice were treated topically 4 times with R848 (n=5, grey bars) or acetone (n=3 white bars) on the right ear. 24 hours after the last treatment mice were anaesthetised and the contralateral ear was visualised using intravital microscopy. Representative images are shown of the contralateral ears (red= dextran, green= CX3CR1-GFP). Non-classical monocytes are quantified as intravascular GFP+ cells per minute (left panel). B) C57BL/6 mice were treated topically 4 times with 100μg of R848. BM-sorted monocytes (4 biological replicates per group) were plated in vitro at a concentration of 1×106 per ml and restimulated overnight with 100μg/ml of R848 or 1μg/ml of LPS. Un-stimulated monocytes were used naïve controls. Cytokines in the supernatant were quantified using a Legendplex Mix and expressed as pg/ml. Data representative of 2 independent experiments. Statistical analysis using unpaired t-test to compare 2 groups (A) and two-way ANOVA with Bonferroni’s multiple comparison test for >2 groups (B). Data are the mean ± SEM; only significant p values are indicated; * p<0.05; ** p<0.01; *** p<0.001; **** p<0.0001.

